# Oct4 is a gatekeeper of epithelial identity by regulating cytoskeletal organization in skin keratinocytes

**DOI:** 10.1101/2023.05.05.539557

**Authors:** Elena D. Christofidou, Marios Tomazou, Chrysovalantis Voutouri, Christina Michael, Triantafyllos Stylianopoulos, George M. Spyrou, Katerina Strati

## Abstract

Oct4 is a pioneer transcription factor regulating pluripotency. However, it is poorly known whether Oct4 has an impact on epidermal cells. We generated *OCT4* knockout clonal cell lines using immortalized human skin keratinocytes to identify a functional role for the protein. Here we report that Oct4-deficient cells transitioned into a mesenchymal-like phenotype with enlarged size and shape, exhibited accelerated migratory behavior, decreased adhesion and appeared arrested at G2/M cell cycle checkpoint. Oct4 absence had a profound impact on cortical actin organization, with loss of microfilaments from cell membrane, increased puncta deposition in the cytoplasm and stress fiber formation. E-cadherin, β-catenin and ZO1 were almost absent from cell-cell contacts while fibronectin deposition was markedly increased in ECM. Mapping of the transcriptional and chromatin profiles of Oct4-deficient cells revealed that Oct4 controls the levels of cytoskeletal, ECM and differentiation related genes, whereas epithelial identity is preserved through transcriptional and non-transcriptional mechanisms.

## INTRODUCTION

The role of the transcription factor Oct4 (encoded by *Pou5f1*) in embryonic stem cells is well documented, both in governing pluripotency *in vitro*^1,2^ as well as *in vivo*^3,4^. Several studies showed that Oct4 is also expressed in tumor cells^5–10^, where it regulates carcinogenesis, but its presence in cells of differentiated or normal somatic tissues has been highly controversial. This is mainly due to the detection of Oct4 pseudogenes^11^, low Oct4 expression levels^12^ or even lack of *OCT4* knockout cells. However, recent research sheds light on the critical role of Oct4 in somatic cells by providing evidence that the protein has a functional role. Specifically, Oct4 activation in smooth muscle cells is atheroprotective by controlling the size and composition of atherosclerotic lesions, resulting in increased plaque stability and thicker fibrous cap^13^. Another study showed that *OCT4* knockout in endothelial cells induced endothelial to mesenchymal transition, plaque neovascularization and mitochondrial defects, thus linking Oct4 with the prevention of phenotypic dysfunction^14^.

The role of Oct4 in skin keratinocytes remains enigmatic. Studies have focused on how transient overexpression of Oct4 in skin keratinocytes changes their differentiation pathway towards induced pluripotent stem cells^15–18^. Skin epidermis is a self-renewing stratified tissue whose mechanical properties are mainly determined by the resident keratinocytes^19,20^. Epidermal stratification is initiated by a monolayer of basal keratinocytes which results into an assembled polarized sheet. The spatial cues that drive the process of polarization are cell-extracellular matrix (ECM) and cell-cell contacts^21,22^. Transmembrane receptors of the integrin superfamily are responsible for cell-ECM interactions whereas the cadherin family is utterly associated with the regulation of cell-cell contacts^23–25^. Moreover, the actin cytoskeleton, a network of interconnected molecules in constant remodeling, generates the mechanical forces essential for the coordination and stabilization of adhesive systems and dictates epithelial cell shape and motility^26,27^. Strong cell-cell adhesion is accomplished through indirect dynamic interaction between E-cadherin and bundles of actin microfilaments^28–30^. Below the plasma membrane of fully formed epithelial sheets lies a cortical belt of actin bundles^31^.

Limited lines of evidence link Oct4 with the regulation of actin cytoskeleton in embryonic stem cells. A previous work that used three-dimensional confocal microscopy to study the distribution of the cytoskeleton in live mouse embryonic stem cells showed that actin depolymerization induces Oct4 binding to chromatin sites, suggesting that actin filamentous network negatively regulates Oct4 ability to bind to DNA^32^. On the contrary, vimentin facilitated the preservation of Oct4-chromatin interactions. These results unveiled a crosstalk between the cytoskeleton and nuclear Oct4 and showed that cytoskeletal components can impact nuclear morphology. Zyxin, a protein that concentrates at focal adhesions and orchestrates mechanical regulation of stress fibers^33^ reduces the expression of *POU5F3* (mammalian *POU5F1/OCT4* homologs) family of genes in *Xenopus laevis* embryos. This is a consequence of a Y-box protein-zyxin complex formation, which leads to *POU5F3* mRNA destabilization and degradation^34^. Interestingly, zebrafish *Pou5f1* deficient maternal and zygotic spiel ohne grenzen mutant embryos develop severe gastrulation defects, which are associated with severe delay in epiboly progression of all three embryonic lineages. Specifically, *Pou5f1* had an impact on cytoskeletal organization and behavior of the enveloping layer and yolk syncytial layer, affected the integrity of the large microtubule arrays located within the yolk syncytial layer and yolk cytoplasmic layer, and delayed the characteristic coordinated epithelial cell shape change of the marginal cells of the enveloping layer^35^. Naturally stored actin in an oocyte nucleus of a *Xenopus* is involved in gene regulation and reprogramming, by transcriptional reactivation of *OCT4*. This signaling process is mediated through Toca-1 protein which enhances *OCT4* reactivation by regulating nuclear actin polymerization^36^. To date, there is no evidence for a functional role for Oct4 within epidermal keratinocytes.

Using CRISPR-Cas9 gene editing technology we have generated *OCT4* knockout clonal cell lines in human immortalized skin keratinocytes (HaCaT). We present evidence that Oct4 preserves cell shape and size, and controls migratory behavior and cell cycle of skin keratinocytes. Oct4 acts as a cytoskeletal organizer, since *OCT4* knockout cells lost the expression of cortical filamentous actin and formed stress fibers. This occurred concomitant with a loss of plasma membrane E-cadherin and ZO1, reduce β-catenin signal but increase fibronectin deposition, suggesting that Oct4 controls the organization of cytoskeletal and ECM landscape. Genes associated with adherens and tight junctions, actin cytoskeleton and ECM were deregulated at the mRNA level. The experimental results are supported by RNA-seq and ATAC-seq data which demonstrate that Oct4 has an extensive influence on cytoskeleton, cell adhesion, cell differentiation and cell behavior. Furthermore, we show that the epithelial phenotype is rescued by an Oct4 mutant protein that lost its DNA binding ability, whilst the transcriptional output of *WASP*, *CDC42* and *ROCK1* are dependent on the transcriptional function of Oct4. These findings provide the first direct evidence that a low-abundance protein, Oct4, has a protective role in skin keratinocytes and a direct involvement with the organization of actin cytoskeleton.

## RESULTS

### Oct4 is a crucial regulator of keratinocyte identity by governing cell size and morphology

We used publicly available data derived from the Protein Atlas to assess the transcript levels of Oct4 in various tissues. Oct4 has a moderate expression in skin compared to other tissues such as kidney and pancreas where it is highly expressed (Figure 1A). To further examine whether Oct4 is specifically expressed in skin keratinocytes we have reanalyzed single cell RNA-seq data from Protein Atlas showing that Oct4 mRNA is high in basal and suprabasal keratinocytes compared to other skin cell types (Figure 1B). These findings are consistent with our experimental data which demonstrate that Oct4 transcripts have been detected and are relatively higher in HaCaT keratinocytes compared to cancer cells HeLa and C33a (Figure 1C). On the contrary, Oct4 protein in HaCaT is undetectable by western blot analysis. To circumvent expression issues, we have used pcDNAOCT4 expression plasmid as a positive control (Figure 1D)^37^.

**Figure 1.**
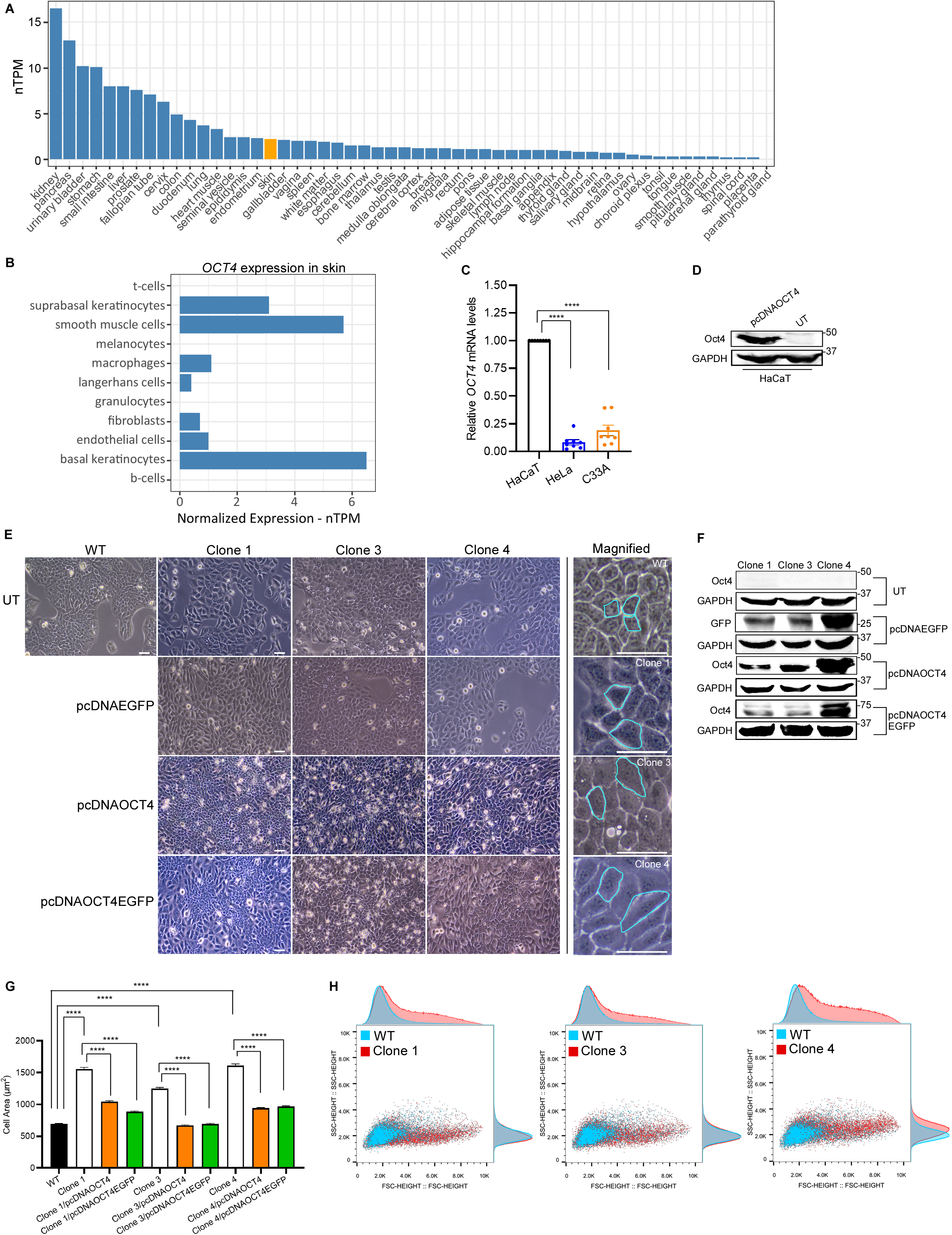
Oct4 controls cell size and morphology of skin keratinocytes. (A) Oct4 transcript levels in different tissues. Data were derived from GTEx dataset available from the Human Protein Atlas (HPA)^72^ (Human Protein Atlas proteinatlas.org). (B) Oct4 expression levels (nTPM) in different skin cell types. Data were derived from single cell RNA-seq dataset available from the HPA^73^. (C) qRT-PCR quantification of Oct4 transcript levels in HaCaT keratinocytes compared to HeLa and C33a cancer cells. n=8, one-way ANOVA. (D) Immunoblot showing Oct4 protein expression in HaCaT cells transfected with pcDNAOCT4 expression plasmid and in untransfected (UT) cells. (E) Representative phase contrast images indicating changes in cell size and morphology. Untransfected or transfected *OCT4* knockout clones with pcDNAEGFP (control), pcDNAOCT4 or pcDNAOCT4EGFP plasmids are compared to WT keratinocytes. Cell size and shape differences of untransfected cells are indicated in magnified images by light blue. Scale bar, 100 μm. (F) Western blot analysis. HaCaT *OCT4* knockout clonal cell lines either untransfected or transfected with pcDNAEGFP, pcDNAOCT4 or pcDNAOCT4EGFP plasmids for 48 h. (G) Quantification of cell size comparing WT keratinocytes to untransfected or transfected *OCT4* knockout clones. n=3, one-way ANOVA. (H) Flow cytometry analysis. Overlayed density plots of WT keratinocytes and *OCT4* knockout clones indicating the shift in cell size (FSC) and complexity (SSC). n=3. See also Figures S1, S2 and Tables S1, S2.

To investigate the functional role of Oct4 in HaCaT skin keratinocytes we knocked out the protein by CRISPR-Cas9 gene editing technology. We generated *OCT4* knockout clonal cell lines (clone 1, -3, and -4) derived from a single cell, which were initially validated by T7E1 assay (Figure S1A). Then, all clones were genotyped by sanger sequencing in combination with Inference of CRISPR Edits (ICE) software for the identification of the type of indels (Figure S1B). We further confirmed genetic knockouts by amplicon sequencing and observed different type of indel formation in each clone (Figure S1C and Table S1). Due to Oct4 low protein abundance in HaCaT we immunoprecipitated Oct4 in cells transfected with pcDNAOCT4 plasmid (positive control), in untransfected WT cells (endogenous Oct4) and in all three *OCT4* knockout clones to assess protein levels (Figure S1D). The results revealed that Oct4 protein was knocked out in all clones.

*OCT4* knockout cells detached more easily from tissue culture plates suggesting that the protein has an impact on cell adhesion (Figure S1E). Notably, loss of Oct4 impacted cell phenotype by inducing a transition from epithelial into more mesenchymal looking morphology. Clonal cells appeared larger in size and different in shape compared to WT cells (Figure 1E, first horizontal and magnified panels). To prove that the phenotypic changes observed is a consequence of the loss of functional Oct4 we performed a rescue experiment where exogenous Oct4 was added to all clones by transient transfection (Figure 1F). One control (pcDNAEGFP) and two different Oct4 expression plasmids were used, pcDNAOCT4 or pcDNAOCT4EGFP. The latter was generated by fusing Oct4 to EGFP and was localized mainly in the nucleus in all *OCT4* knockout cells (Figure S2A). Exogenously added Oct4 rescued cell phenotype in a subset of cells (Figure 1E, third and fourth horizontal panels) whereas clonal cells transfected with pcDNAEGFP control showed no morphological change (Figure 1E, second horizontal panel). The difference in cell size was quantified by measuring cell area in single cells and found statistically significant and consistent for all comparisons made. Specifically, *OCT4* knockout clones had significantly larger size compared to WT cells, while when transfected with Oct4 expression plasmids clonal cells had a significant reduction in size than their untransfected counterparts (Figure 1G). Moreover, comparing cell size between *OCT4* knockout cells transfected with pcDNAOCT4 or pcDNAOCT4EGFP to WT cells resulted in no statistically significant difference (Figure S2B) suggesting that Oct4 re-expression restored the epithelial phenotype.

Furthermore, we evaluated the impact of *OCT4* knockout on keratinocyte cell size and distribution by flow cytometry. Clonal cells shifted towards higher forward scatter values indicating that are larger in size compared to WT (Figure S2C). Image overlay of WT cells with clonal cell lines clearly shows the change in cell size and complexity, and the transition into a different type of cell (Figure 1H).

These data indicate that Oct4 is responsible for maintaining the epithelial characteristics and that phenotypes observed are unlikely to represent off-target effects of gene editing.

### Keratinocyte Oct4 deficiency accelerates migratory behavior and inhibits cell cycle progression

Keratinocyte migration is an essential step for re-epithelialization during the vital process of wound healing. This is essential for skin barrier restoration^38^. Another important activity is keratinocyte migration from the basal layer toward the surface of the skin during differentiation^39^. We investigated whether *OCT4* knockout in HaCaT impacts cell migration through wound healing assays. Cell motility was examined with two different set ups. Firstly, an end-point/conventional wound healing assay was used, where images were taken at timepoint zero and 20 hours after wounding. Secondly, a live time-lapse imaging approach was used with 2 hours intervals between each acquisition timepoint for a total of 20 hours. The results showed that knockout of *OCT4* accelerated keratinocyte migration and wound healing gap closure (Figure 2A and Movies S1-S4). This observation was consistent for all clonal cell lines (clones 1, -3 and -4) with the percentage of gap closure being significantly larger for the clones compared to WT cells (Figure 2B). Interestingly, the increased migratory capacity of clonal cells was not accompanied by accelerated proliferation. Cell numbers were measured at 24, 48 and 72 hours after seeding. Proliferation was unaltered for 24 and 48 hours for all conditions, while WT and *OCT4* knockout cells showed increased growth rate at 72 hours (Figure 2C).

**Figure 2.**
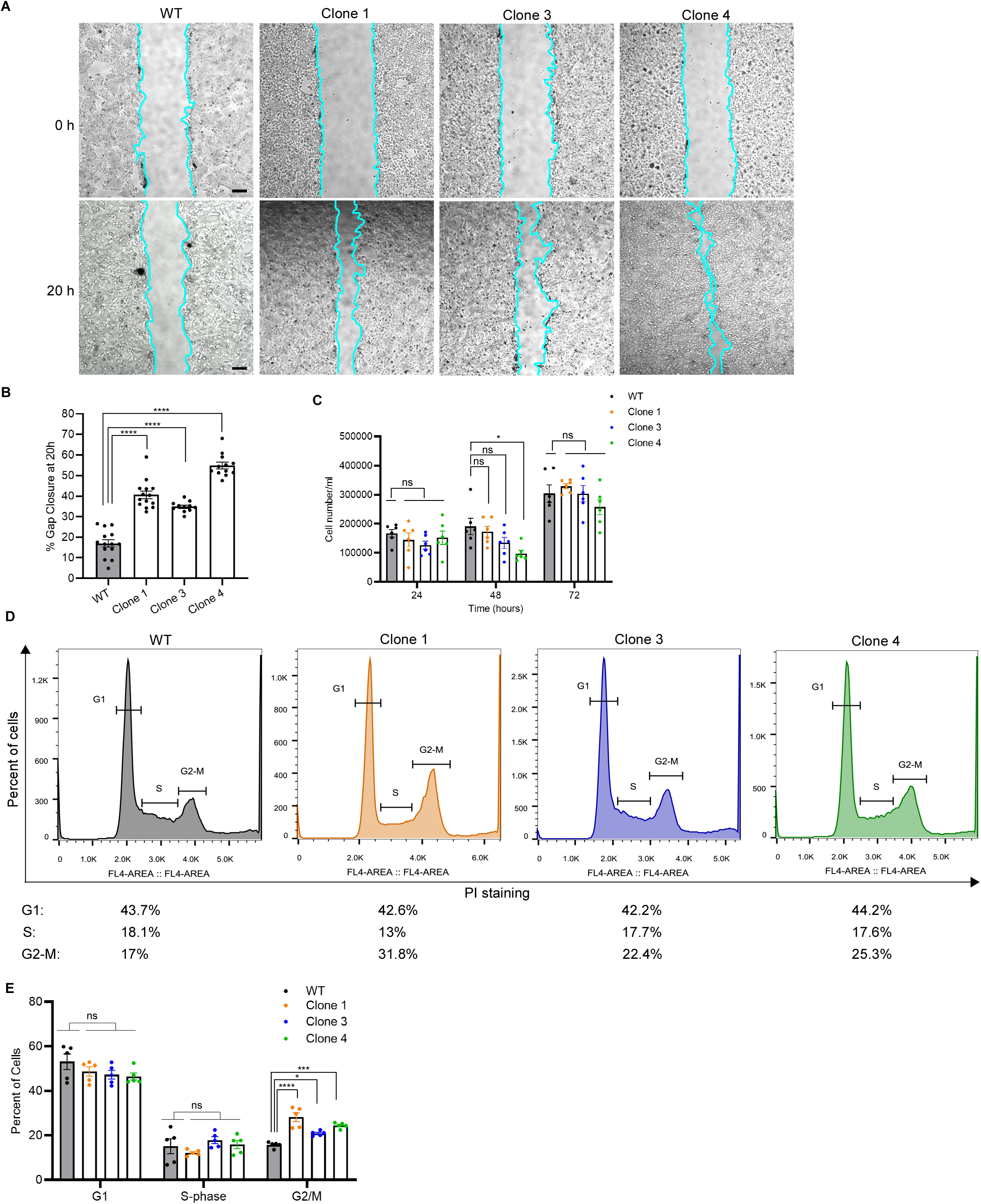
Oct4 regulates migratory behavior and cell cycle of epidermal keratinocytes. (A) Representative images of wound healing assay of WT and *OCT4* knockout cells from 0 h to 20 h. Scale bar, 600 μm. n=2 for end-point and n=2 for live cell migration experiments. (B) Quantitatively analysis of wound healing assay in (A) by AxioVision 4.8 software. n=4, one-way ANOVA. (C) Effect of *OCT4* knockout on cell proliferation at 24, 48 and 72 h. WT keratinocytes compared to *OCT4* knockout clones. n=3, one-way ANOVA. (D) Histograms demonstrating cell cycle distribution of WT keratinocytes and *OCT4* knockout clonal cells after PI staining. (E) Quantitatively analysis of WT keratinocytes and *OCT4* knockout clones at G1, S and G2/M phases of the cell cycle. n=5, one-way ANOVA. See also Movies S1-S4 and Figure S3.

Proper execution of cell division is associated with the control of cell cycle checkpoints. Oct4 promotes the progression of the cell cycle in embryonic stem cells by stimulating the entry from G1 to S-phase^40^. Therefore, we examined whether Oct4 regulates cell cycle progression in HaCaT keratinocytes. WT and *OCT4* knockout cells were stained with propidium iodine (PI) and analyzed by flow cytometry. Loss of Oct4 inhibited G2/M cell cycle progression with clonal cells arrested at this checkpoint (Figure 2D). The percentage of *OCT4* knockout cells arrested in G2/M phase was significantly larger compared to WT cells, while their difference in G1 or S-phase was not statistically significant (Figure 2E). G2/M checkpoint blocks the entry into mitosis when DNA is damaged or when DNA replication is incomplete, suggesting that Oct4 might be involved in mechanisms that regulate DNA damage repair.

We further compared the distribution of WT keratinocytes and clonal cells in G1, S and G2/M cell cycle phases (Figure S3A). *OCT4* knockout clonal cells, with the majority larger in size than WT cells (Figure 1G), were equally distributed in the three cell cycle phases (Figure S3B) suggesting that their mesenchymal-like phenotype accompanied by a larger size was a consequence of loss of functional Oct4 and not due to arrest at G2/M checkpoint.

The above data demonstrate a functional role for Oct4 in skin keratinocytes by controlling cell migration and cell cycle progression.

### Loss of Oct4 in skin keratinocytes promotes rearrangements in actin cytoskeleton and affects fibronectin deposition

Loss of Oct4 results in changes in morphology and size of skin keratinocytes and Oct4 restoration in the Oct4-deficient cells reversed this process. Given these results and the known crosstalk between Oct4 and cytoskeleton in embryonic stem cells^32^, we hypothesized that Oct4 induces these alterations via organization of actin cytoskeleton. To evaluate this hypothesis, we used confocal microscopy to identify changes in actin microfilament structures. To visualize the architecture of the F-actin cytoskeleton we stained WT and Oct4-deficient cells with phalloidin on glass coverslips. We noticed a dramatic change in actin bundles organization with *OCT4* knockout cells loosing cortical actin from plasma membrane and cell-cell boundaries (Figure 3A). In addition, F-actin cortex appeared disorganized and in the form of puncta. Higher magnification images revealed the formation of stress fibers in *OCT4* knockout clones and loose meshwork of actin filaments (Figures 3B and S4A) suggesting that cells acquired a mesenchymal phenotype. Firstly, we quantified the mean fluorescence intensity to examine F-actin concentration. The average intensity of F-actin is significantly decreased in clonal cells compared to WT (Figure S4B) To identify whether actin is reorganized from the plasma membrane to the cytoplasm we measured the ratio of the membrane mean fluorescence to the cytoplasmic mean fluorescence. Cortical actin membrane/cytoplasm ratio is significantly decreased in *OCT4* knockout cells compared to WT (Figure 3C), suggesting that Oct4 absence induced actin rearrangements in keratinocytes and specifically from the cell membrane to the cytoplasm. To gain more insight into the distribution of cortical actin we specifically quantified fluorescence intensity on the cell membrane and in the cytoplasm. In WT cells, fluorescence was localized on the membrane while in Oct4-deficient cells was diminished (Figure 3D). Intensity variations are also observed in intensity profiles, where the tracks clearly show that cortical actin is lost from the cell membrane of clonal cells (Figure S4C). The opposite occurred in the cytoplasm where we observed increased intensity in clonal cells compared to WT (Figure 3E) indicating that actin-containing cellular features, in the form of stress fibers or puncta, were predominantly concentrated in the cytoplasm.

**Figure 3.**
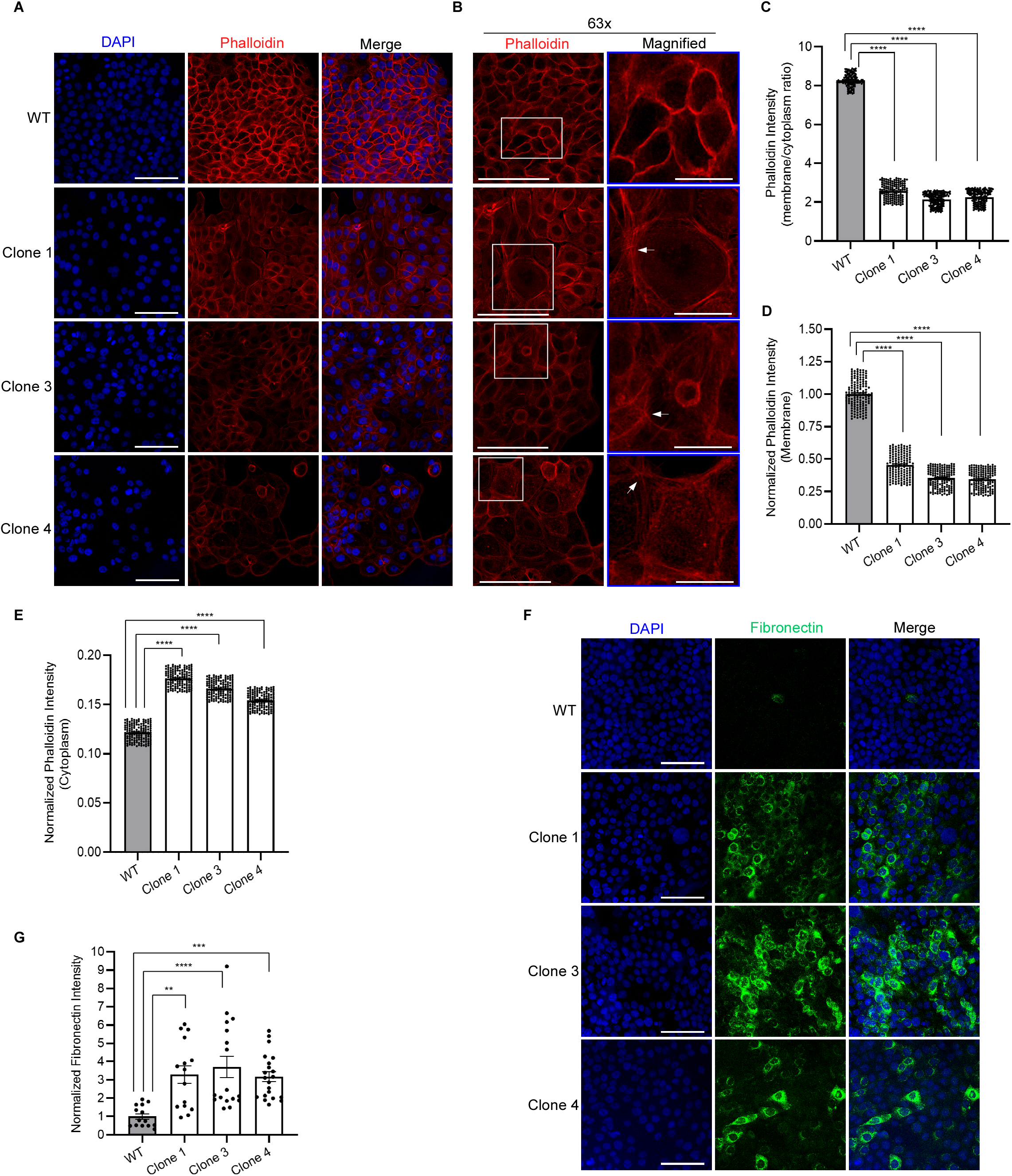
Oct4 impacts on actin cytoskeleton organization and fibronectin deposition in ECM. (A) Representative confocal images of WT keratinocytes and *OCT4* knockout clones stained with phalloidin for F-actin visualization at 40x magnification. Scale bars, 100 μm. (B) Representative images of WT keratinocytes and *OCT4* knockout clones stained with phalloidin at 63x magnification. Magnified area (white rectangle): detailed visualization of stress fibers. White arrows point to stress fibers. Scale bars, 100 μm and 25 μm for magnified images. (C) Graph shows the ratio of membrane/cytoplasm of phalloidin fluorescence intensity distribution comparing WT keratinocytes to *OCT4* knockout clones. n=3, one-way ANOVA. (D) Quantification of phalloidin localized signal of fluorescence on the plasma membrane. n=3, one-way ANOVA. (E) Quantification of phalloidin fluorescence intensity specifically in the cytoplasm. n=3, one-way ANOVA. (F) Representative confocal images of WT keratinocytes and *OCT4* knockout clones stained with fibronectin. Scale bars, 100 μm. (G) Quantification of fibronectin fluorescence intensity showing its deposition in ECM. n=3, one-way ANOVA. See also Figure S4.

Loose cell adhesions in Oct4-deficient cells point to dysregulated and altered ECM. Basal keratinocytes of the dermal-epidermal junction lie on a specialized and mechanically supportive network of ECM proteins^41^. Fibronectin is a crucial factor for enhanced motility and spreading of keratinocytes^42^. To elucidate the role of Oct4 in the regulation of ECM components we stained WT and *OCT4* knockout cells with fibronectin on glass coverslips. Imaging revealed that fibronectin deposition in ECM was highly increased in the clones (Figure 3F), while quantification of mean fluorescence intensity showed a significant difference between WT and clonal cell lines (Figure 3G).

Our data demonstrate a strong influence of Oct4 in the regulation of cortical actin organization and in the control of endogenous fibronectin secretion in ECM.

### Oct4 preserves the formation of adherens and tight junctions in skin keratinocytes

The primary contact points between cells and their environment are cellular adhesions, with cadherin complex being the main mediator of cell-cell adhesion in all soft tissues. Cadherin-catenin interactions are responsible for organizing actomyosin by tethering to microfilaments, thus maintaining cell adhesive properties, polarity and integrity^31^. We thus went on to examine if loss of adhesion of Oct4-deficient cells is associated to E-cadherin or β-catenin impairments. We show that E-cadherin and β-catenin are almost completely lost from the cell-cell contacts of *OCT4* knockout cells, whereas both proteins are mainly localized at the plasma membrane of WT keratinocytes (Figure 4A and S5A). These observations are consistent for all clonal cell lines, suggesting that even though different indels are formed in each clone the final phenotypic outcome is the same for all clones. In higher magnification images we indicate that in WT cells E-cadherin is colocalized with β-catenin at the cell membrane, thus contributing to the cell-cell contact maintenance, while in Oct4-deficient cells this colocalization is lost with cells appearing more disorganized and with no epithelial identity (Figure 4B). Quantification of their spatial overlap showed that colocalization was significantly attenuated in clones as opposed to WT cells (Figure S5B). We next determined protein concentration by measuring the mean fluorescence intensity for β-catenin and E-cadherin, where in both cases signal was significantly reduced in clonal cells compared to WT (Figures S5C and S5D). To gain more insight into E-cadherin and β-catenin spatial distribution and localization we have measured the ratio membrane/cytoplasm and the fluorescence intensity specifically on the cell membrane. Again, for both proteins the ratio and membrane intensity were significantly decreased in Oct4-deficient cells, suggesting reorganization and disassembly of adherens junctions, confirming transition into a mesenchymal-like phenotype and indicating that loss of Oct4 affects E-cadherin/β-catenin interaction and localization (Figures 4C, 4D, 4E and 4F).

**Figure 4.**
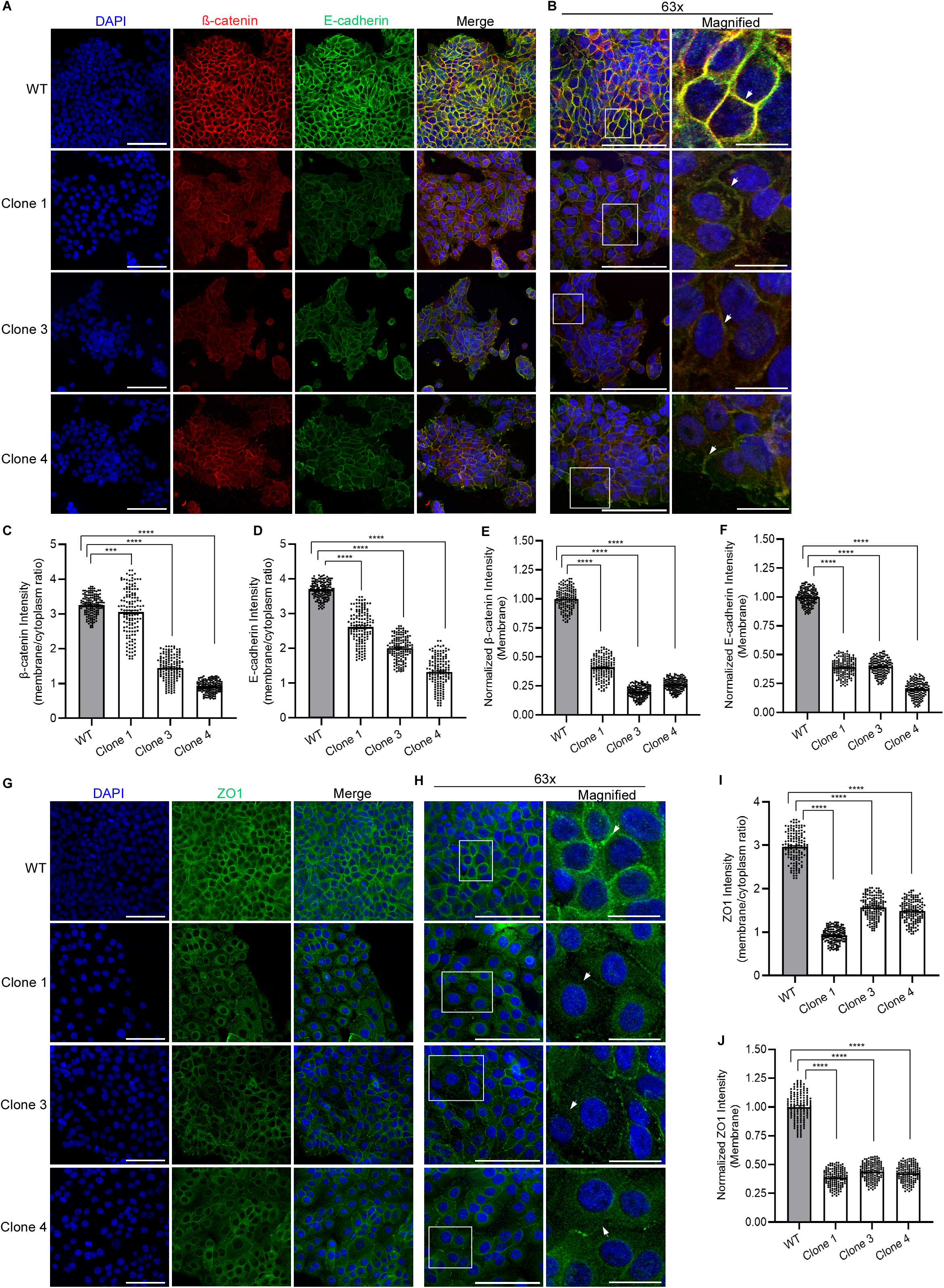
Oct4 stabilizes the formation of adherens and tight junctions at cell-cell contacts and guides E-cadherin and β-catenin localization. (A) Representative confocal images of WT keratinocytes and *OCT4* knockout clones indicating the distribution of β-catenin and E-cadherin at cell-cell contacts using 40x objective. n=3, scale bars, 100 μm. (B) Representative merged images at 63x magnification and magnified area (white rectangle), showing localization of β-catenin and E-cadherin (white arrows). Scale bars, 100 μm and 25 μm for magnified images. (C) Graph shows the ratio of membrane/cytoplasm of β-catenin fluorescence intensity distribution comparing WT keratinocytes to Oct4-deficient clones. n=3, one-way ANOVA. (D) Graph indicates the ratio of membrane/cytoplasm of E-cadherin fluorescence intensity distribution comparing WT keratinocytes to *OCT4* knockout clones. n=3, one-way ANOVA. (E) Quantification of β-catenin fluorescence intensity specifically on the plasma membrane. n=3, one-way ANOVA. (F) Quantification of E-cadherin localized signal of fluorescence at the cell membrane n=3, one-way ANOVA. (G) Representative confocal images of WT keratinocytes and *OCT4* knockout clones demonstrating the localization of ZO1 at cell-cell contacts using 40x objective. n=3, scale bars, 100 μm. (H) Representative images at 63x magnification and magnified area (white rectangle), showing detailed localization of ZO1 (white arrows). Scale bars, 100 μm and 25 μm for magnified images. (I) Graph indicates the ratio of membrane/cytoplasm of ZO1 fluorescence intensity distribution comparing WT keratinocytes to *OCT4* knockout clones. n=3, one-way ANOVA. (J) Quantification of ZO1 fluorescence intensity specifically at the plasma membrane n=3, one-way ANOVA. See also Figure S5.

Tight junctions are the most apical elements of the junctional complex, and they are crucial for the regulation of epidermal barrier by controlling the movement of ions and solutes in between keratinocytes^43^. To identify whether Oct4 is associated with the regulation of tight junctions in skin keratinocytes we stained WT and Oct4-deficient cells with the tight junction scaffolding protein ZO1 on glass coverslips. Confocal imaging showed that ZO1 was diminished from the cell membrane area of *OCT4* knockout cells while we observed that cells were larger and irregular in size (Figure 4G and S5G). Magnified images indicate that ZO1 was almost absent from the tight junctions of Oct4-deficient cells (Figure 4H) having as a consequence loss of epithelial cell-cell contacts. These observations were again consistent in all clonal cell lines. In addition, the mean fluorescence intensity of clonal cells, indicating protein concentration, was significantly reduced compared to WT cells (Figure S5H). Since ZO1 is a junctional adaptor protein we explored whether its localization and distribution change when Oct4 is absent. The membrane/cytoplasm ratio is significantly attenuated in clonal cells compared to WT keratinocytes (Figure 4I), whereas quantification of the signal specifically on the plasma membrane and evaluation of intensity distribution along the cell diameter of single cells show similar results (Figures 4J and S5I).

Overall, these findings show that Oct4 guides the spatial distribution of E-cadherin, β-catenin and ZO1 in skin keratinocytes which subsequently leads to the preservation of adherens and tights junctions. Furthermore, Oct4 impacts on the colocalization of E-cadherin with β-catenin, suggesting that the protein might enhance the interaction between them and possibly influence mechanisms associated to Wnt signaling.

### Oct4 governs the transcriptional landscape of cytoskeletal and ECM genes

A unique, stable, and self-sufficient gene expression pattern characterizes epithelial identity. Transition from one cell type to another is accompanied by alterations in gene expression. The disorganizing effect of Oct4 deficiency on keratinocyte cytoskeleton and ECM led us to examine whether Oct4 alters the transcript levels of genes associated with the regulation of actin cytoskeleton. We first tested genes involved in the formation of adherens and tight junctions. E-cadherin and ZO1 mRNA levels were significantly increased in *OCT4* knockout cells compared to WT, while β-catenin transcripts were significantly decreased in all clonal cell lines (Figure 5A). E-cadherin and ZO1 increased mRNA levels in clones is opposite to their decreased localized fluorescence signal mentioned previously, suggesting that Oct4 might control protein localization and gene expression through a different mechanism.

**Figure 5.**
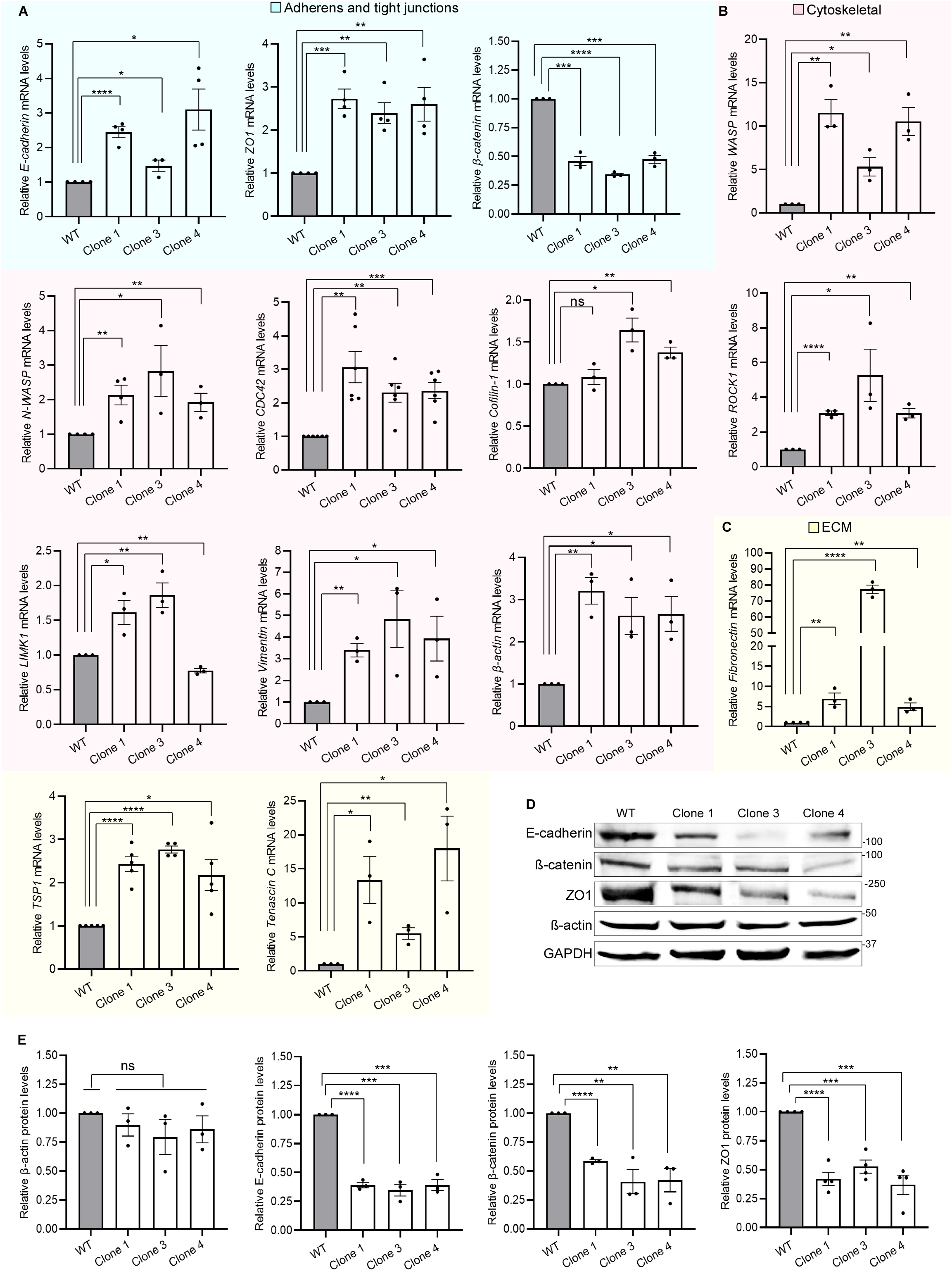
Oct4 determines the transcriptional landscape of cytoskeletal, ECM and adherens and tight junction genes in skin keratinocytes. (A) qRT-PCR quantification of adherens and tight junction genes. Comparison of gene expression between WT skin keratinocytes and *OCT4* knockout clones. n=at least three experiments, one-way ANOVA. (B) Transcript levels of cytoskeletal genes. n=at least three experiments, one-way ANOVA. (C) qRT-PCR quantification of ECM genes. n=at least three experiments, one-way ANOVA. (D) Representative western blot images of E-cadherin, β-catenin, ZO1 and β-actin protein levels in WT and *OCT4* knockout cells. (E) Quantification of β-actin, E-cadherin, β-catenin and ZO1 relative protein expression. n=3, unpaired Student’s t-test. See also Table S2.

Next, we tested genes which interact with the actin cortex and regulate its polymerization. Actin polymerization is a prerequisite for the formation of lamellipodia and filopodia from the cell surface. Specifically, CDC42 induces actin polymerization by stimulating a family of proteins called Wiskott-Aldrich Syndrome proteins (WASP) with main members WASP and N-WASP^44^. Transcript levels of CDC42, WASP and N-WASP were significantly increased in Oct4-deficient cells compared to WT (Figure 5B). ROCK1 an upstream regulator of LIMK1, which in turn controls Cofilin1 phosphorylation, is a well-characterized effector of the small GTPase RhoA^45^. All proteins have important roles in actin polymerization, thus we investigated how Oct4 affects their transcriptional output. Cofilin1 mRNA levels were increased only in clones 3 and 4, whereas LIMK1 was increased in clones 1 and 3 but decreased in clone 4. This discrepancy between clones might be attributed to the different indel formation in each clone or to single nucleotide mutations unique to each clone generated after a short period in culture^46^. However, ROCK1 transcript levels were significantly increased (at least 3-fold change) in *OCT4* knockout cells compared to WT (Figure 5B). Vimentin, a major protein of the intermediate filament family usually found in mesenchymal cells but not in keratinocytes^47^, was upregulated in all Oct4-deficient cells. Similar results were obtained for β-actin transcript levels (Figure 5B).

Structural support for the cells and tissues is provided by a dynamic 3-dimensional network of macromolecules known as ECM. Fibronectin, tenascin C and thrombospondin 1 (TSP1) are ECM proteins which support keratinocyte interactions during wound healing^48,49^ and provide an environment for cell adhesion^50^. Transcript levels of fibronectin in Oct4-deficient clonal cell lines were markedly increased compared to WT cells (Figure 5C), and this is in agreement with its increased deposition in ECM as mentioned previously. Similarly, TSP1 and tenascin C were also upregulated in all clones examined (Figure 5C).

Relative protein levels of β-actin remained unchanged in all conditions tested (Figures 5D and 5E), even though its cellular localization was dramatically altered in Oct4-deficient cells. On the contrary, E-cadherin, β-catenin and ZO1 protein expression was significantly decreased in all clonal cell lines compared to WT (Figures 5D and 5E), which agrees with their reduced fluorescence intensity observed previously.

Discrepancies between transcript and protein expression levels of either E-cadherin or ZO1 might be attributed to post-transcriptional mechanisms which can shape protein levels independently of mRNA abundance. Moreover, translation rates which are influenced by mRNA sequence and a protein’s half-life that affects protein concentration independent of mRNA levels are important factors that might contribute to differences between cellular protein levels and transcripts^51^. Overall, the above findings prove that the majority of the results are consistent among clonal cells and that Oct4 maintains the transcriptional network governing cytoskeletal and ECM genes.

### Oct4 loss of function reshapes transcriptional and chromatin landscapes

To profile the transcriptomic consequences of Oct4 loss, we performed RNA-seq in WT keratinocytes, clone 1 and clone 4 knockout cell lines. Two replicate samples from each condition were sequenced. A heatmap of sample-to-sample distances calculated using the full transcript vector (Figure S6A) shows a close relationship between sample replicates but greater distances between each of the clones and the WT. From the differential expression analysis we obtained a total of 2135 and 1830 significantly differentially expressed genes (DEGs) in clone 1 and clone 4, respectively, compared to WT condition, of which 806 genes were common to both comparisons (Figure 6A). The dominant transcriptional alteration was the deregulation of genes involved in cytoskeletal and matrix organization and cell differentiation. Specifically, genes associated with cell adhesion, actin cytoskeleton remodeling, migration, matrix deposition and cell cycle progression were either upregulated (*CCDC80*, *KRT7*, *TIMP2*, *PODXL*, *ROBO1*, *KRT8*, *COL4A2*) or downregulated (*DKK3*, *SPRR1B*, *CCND2*, *SOX2*, *CORO6*, *LAMB1*) when Oct4 was absent (Figure 6B). Some of these DEGs were common to both clones (*FN1*, *ROBO1*, *PODXL*) and some were unique (*KRT7*, *COL4A2*). These results show that Oct4 has crucial roles in the regulation of cytoskeletal architecture in skin keratinocytes and confirm our experimental findings demonstrated previously.

**Figure 6.**
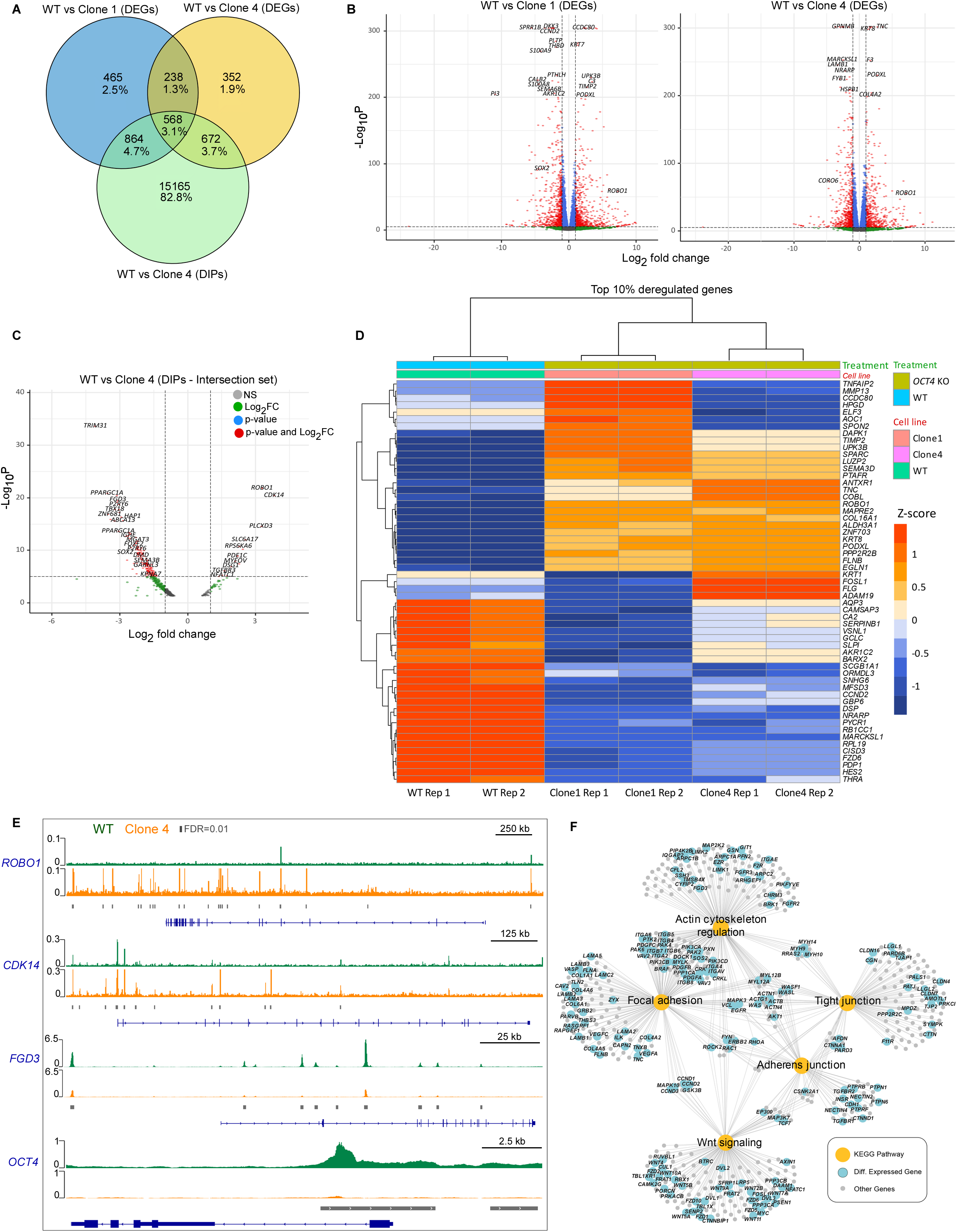
Loss of Oct4 correlates with alterations in transcriptome and chromatin accessibility by deregulating cytoskeletal genes. (A) Venn diagram indicating the unique and common DEGs in the two RNA-seq comparisons shown in (B) and the DIPs obtained from the ATAC-seq comparison (WT vs clone 4). (B) Volcano plots showing the significance (-Log10P) versus the magnitude (Log2FC) of the DEGs comparing WT to clone 1 (left) or to clone 4 (right). (C) Volcano plot indicating the DIPs derived from the intersection dataset. Intersection set: genes found both differentially expressed in the RNA-seq datasets (WT compared to clone 1 or clone 4) and their ATAC-seq peaks (WT compared to clone 4) that overlap their genomic positions. (D) Heatmap and hierarchical clustering of the significant top 10% DEGs derived from the intersection dataset. *OCT4* knockout clones 1 and 4 show distinct clusters of genes that are up- or downregulated compared to WT samples. (E) IGV visualized peak traces for *ROBO1, CDK14, FGD3* and *OCT4* genes comparing WT (green) to clone 4 (orange). Grey lines correspond to peaks which are significantly changed between WT and clone 4. (F) Gene to pathway bipartite network showing the unique and common membership of genes in pathways of interest enriched from KEGG. Yellow nodes correspond to pathways, blue nodes to significantly DEGs (intersection dataset) and grey nodes to genes not significantly perturbed. See also Figures S6 and S7.

Oct4 has the ability to bind to a large number of motifs in inaccessible regions of chromatin, during induced pluripotent stem cell reprogramming^52,53^. To characterize the effects of *OCT4* knockout in genome-wide chromatin accessibility we harvested cells from WT and clone 4 conditions to perform Assay for Transposase-Accessible Chromatin with high-throughput sequencing (ATAC-seq)^54^. We have identified 15165 significant differentially intense peaks (DIPs). The majority of the peaks were present at intronic (33%) or distal intergenic regions (28%), while a large subset of peaks (36%) corresponded to promoters and UTRs (Figure S6B). Analysis of both RNA-seq and ATAC-seq data revealed that 1432 or 1240 genes found deregulated in clone 1 or clone 4 RNA-seq datasets, respectively, had significant DIPs overlapping their genomic positions (Figure 6A). The integration of the three datasets (two DEG and one DIP sets) resulted in 568 deregulated genes (Figure 6A). This was defined as the intersection set and includes the genes found differentially expressed in both the RNA-seq datasets and have differentially intense ATAC-seq peaks overlapping their genomic positions. Nearly 71% of these genes had a DIP overlapping their transcriptional start site (TSS). Notably, among these DIPs (intersection set) we have identified *ROBO1*, *CDK14*, *DSG1* and *TGFBR3* genes to have regions with open chromatin in Oct4-depleted cells, while *FGD3*, *SOX2*, *TBX18* and *P2YR6* genes to possess close chromatin structure (Figure 6C). These observations are in agreement with our functional data which show that Oct4 regulates cell migration, cell cycle and adhesion and actin microfilament remodeling by reshaping the chromatin landscape of genes involved in these processes.

Furthermore, to identify DEG patterns that can differentiate robustly the WT and the *OCT4* knockout clones we selected the top 10% most abundant significantly deregulated genes from the intersection set and performed hierarchical clustering (Figure 6D). Most of the selected genes show similar expression levels with the exception of a small anticorrelated cluster between knockout clones. Expectantly, we obtained a clear cluster separation showing a similar profile between replicates and clones, and a wider distance of each of the clones with the WT. Nearly all selected genes are associated with actin cytoskeleton, cell adhesion, cell differentiation, migration and ECM organization suggesting that Oct4 control epigenetic-related mechanisms utterly involved in these signaling pathways.

ROBO1, the receptor for SLIT proteins is important for axonal guidance^55^ and for dynamic tuning of cell-cell and cell-matrix interactions^56^. CDK14 is a member of the atypical cyclin dependent kinases family and is implicated in the regulation of cell cycle by binding to G2/M cyclins, cyclin Y and cyclin Y-like 1^57^. We show that loss of Oct4 in HaCaT cells dramatically increases chromatin accessibility of both genes, while the opposite is observed for *FGD3* gene (Figure 6E and S6C), which plays a role in actin cytoskeleton, controls cell shape and promotes the formation of filopodia^58^. Strikingly, loss of Oct4 induced a decrease in chromatin accessibility close to its own 5’-UTR and at promoter region (Figure 6E), suggesting that the gene might regulate its own expression by binding to regulatory elements.

To investigate the functional role of the deregulated genes we performed an overrepresentation analysis (ORA) against KEGG pathways using either the intersection set or the individual DEGs and DIPs. Supportive to the other experimental findings, the analysis revealed that among others, focal adhesion, regulation of actin cytoskeleton, tight junction, adherens junction and Wnt signaling pathways were commonly enriched in all three datasets (Figure S6D). 23 and 26 unique pathways were identified for each of the two RNA-seq comparisons: WT vs clone 1 or WT vs clone 4, respectively, and 111 common pathways. Overall, 87 pathways were common for all three datasets examined (Figure S6E). We used the 5 pathways of interest to generate a bipartite gene to pathway network, in order to analyze the gene to pathway membership of the intersection set (Figure 6F). *RhoA* was found to be a member of all pathways.

To better define the role of Oct4 in regulating chromatin accessibility of genes associated with cytoskeletal functions in the 5 pathways of interest, we assessed the distribution of the significant DIPs based on their log2 fold change values (WT vs clone 4). Strikingly, the majority of DIPs of these genes were downregulated in clone 4 (closed chromatin) (Figure S7A), suggesting that the presence of Oct4 in keratinocytes is crucial to ensure a balanced control of cytoskeletal organization. Next, we combined RNA-seq with ATAC-seq data to compare the change in accessibility in promoter proximal peaks to that of the corresponding gene expression levels. 694 genes showed both downregulation of transcript levels and decreased chromatin accessibility whilst 126 genes showed the opposite. Only 4 genes with increase in chromatin accessibility exhibited reduced gene expression (Figure S7B). Impressively, 5332 DIPs were overall downregulated and only 277 DIPs were upregulated (Figure S7C), indicating that Oct4 preferentially maintains chromatin at an open state in human keratinocytes.

In summary, these findings indicated that Oct4 participates in a core cytoskeletal regulatory circuit which drives cortical actin organization by interconnecting it to other pathways that affect epithelial cell shape, adhesion and function.

### Oct4 controls keratinocyte phenotype through non-transcriptional mechanisms

A previous study showed that phosphorylation of Oct4 at T234 and S235, which are located within the Oct4 homeobox domain, negatively regulates the protein by disrupting sequence-specific DNA binding, thus rendering Oct4 transcriptionally inactive^59^. To better understand whether the phenotypic and transcriptional changes that we observed in *OCT4* knockout cells are attributed to Oct4 pioneer function as a transcription factor or to a non-transcriptional level of regulation we used three different Oct4 constructs. pCEP-OCT4-WT which resembles the WT protein, pCEP-OCT4-AA that mimics the non-phosphorylated protein (similar characteristics to WT construct), and pCEP-OCT4-EE which mimics the constitutively phosphorylated Oct4 protein with no DNA binding ability^59^ (Figure 7A).

**Figure 7.**
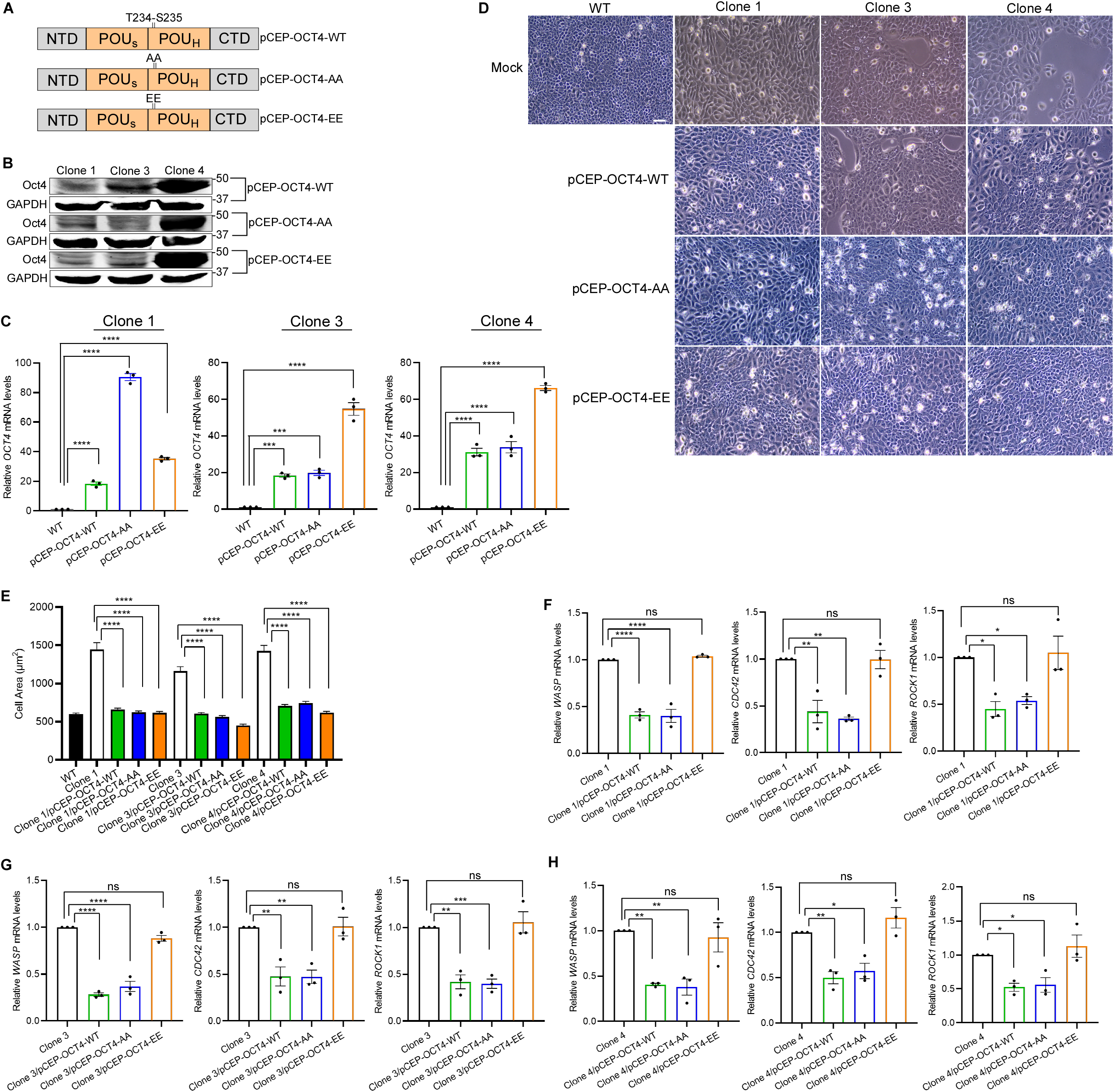
Phenotypic and specific gene expression changes in skin keratinocytes are not utterly dependent on Oct4 transcriptional function. (A) Schematic depicting Oct4 pCEP constructs. pCEP-OCT4-WT (upper), pCEP-OCT4-AA resembles/mimics the WT construct (middle) and pCEP-OCT4-EE mutant does not bind to DNA (lower). (B) Representative western blot images showing Oct4 protein expression in *OCT4* knockout clones transfected with the constructs mentioned in (A). (C) qRT-PCR quantification of *OCT4* mRNA levels comparing WT cells to *OCT4* knockout clones transfected with the constructs mentioned in (A). n=3, one-way ANOVA. (D) Representative phase contrast images demonstrating changes in cell size and morphology. *OCT4* knockout clones untransfected or transfected with pCEP-OCT4-WT, pCEP-OCT4-AA, or pCEP-OCT4-EE constructs compared to WT cells. Scale bar, 100 μm. (E) Quantification of cell size comparing untransfected *OCT4* knockout cells to clones transfected with pCEP-OCT4-WT, pCEP-OCT4-AA, or pCEP-OCT4-EE constructs. n=3, one-way ANOVA. (F) qRT-PCR quantification of *WASP*, *CDC42* and *ROCK1* transcript levels in untransfected clone 1 compared to clone 1 transfected with pCEP-OCT4-WT, pCEP-OCT4-AA, or pCEP-OCT4-EE constructs. n=3, one-way ANOVA. (G) qRT-PCR quantification of *WASP*, *CDC42* and *ROCK1* transcript levels in untransfected clone 3 compared to clone 3 transfected with pCEP-OCT4-WT, pCEP-OCT4-AA, or pCEP-OCT4-EE constructs. (H) qRT-PCR quantification of *WASP*, *CDC42* and *ROCK1* transcript levels in untransfected clone 4 compared to clone 4 transfected with pCEP-OCT4-WT, pCEP-OCT4-AA, or pCEP-OCT4-EE constructs. See also Table S2.

Oct4-deficient cells were transiently transfected with each of these constructs and Oct4 protein and mRNA levels were tested by western blot (Figure 7B) and qRT-PCR (Figure 7C), respectively. Oct4 protein was detected in all clones after 48h of transfection and Oct4 transcript levels were increased at least 20-fold compared to WT sample. *OCT4* knockout cells transfected with pCEP-OCT4-WT or pCEP-OCT4-AA mutant restored keratinocyte phenotype as observed with phase-contrast microscopy (Figure 7D). Remarkably, Oct4 clones transfected with the constitutively phosphorylated pCEP-OCT4-EE mutant, previously shown to be deficient for transcriptional activity and inducing reprogramming, also transitioned back to a WT-like phenotype (Figure 7D), suggesting that functions of Oct4 not strictly limited to its transcriptional role may regulate epithelial identity. The changes in cell size were quantified by measuring cell area. Oct4-deficient cells transfected with pCEP constructs were significantly smaller in size compared to their mock-transfected counterparts (Figure 7E).

Loss of Oct4 impacted the transcriptional profile of cytoskeletal and ECM genes, as we proved by qRT-PCR and RNA-seq. We hypothesized that some of these changes might be related to a transcriptional Oct4 role. Indeed, we observed that *WASP*, *CDC42* and *ROCK1* transcripts in *OCT4* knockout cells transfected with pCEP-OCT4-WT or pCEP-OCT4-AA mutant were significantly decreased compared to mock transfected clones, whereas in clonal cells transfected with the pCEP-OCT4-EE mutant mRNA levels remained unaltered and not significant (Figure 7F, 7G and 7H). This result was consistent in all clones examined.

Overall, these results suggest that restoration of epithelial phenotype in these clonal cell lines might be attributed in part to a non-transcriptional Oct4 mechanism, which also serves a crucial transcriptional regulatory role in skin keratinocytes.

## DISCUSSION

Oct4 is a pioneer transcription factor fundamental for maintaining embryonic stem cells in undifferentiated state^3^. The protein exerts most of its activities by recognizing and binding its target sequences in inaccessible chromatin, thus establishing new transcriptional networks^60^. The main notion until recently was that Oct4 is completely inactive and non-functional in somatic cells, possibly due to its low expression levels. However, previous work showed that Oct4 is implicated in the regulation of atherosclerosis in vascular smooth muscle cells^13^ and in endothelial cells^14^. To our knowledge the current study is the first demonstrating an important functional role for Oct4 in skin keratinocytes, even though its protein levels are almost undetectable. Loss of Oct4 function, through generation of *OCT4* knockout clonal cell lines, induced epithelial cell transitioning into a mesenchymal-like phenotype with crucial impact on cell adhesion, size and shape. Ectopic expression of WT Oct4 in Oct4-deficient cells can rescue the keratinocyte phenotype by restoring epithelial characteristics. These findings suggest that fine tuning of Oct4 levels is necessary for preserving epithelial identity, despite its low protein abundance.

Oct4 expression is indispensable for the generation of induced pluripotent stem cell lines^61^, which are vital for tissue engineering during the wound healing process. Here we show that Oct4 has a profound impact on migratory behavior of skin keratinocytes. Knockout of Oct4 accelerated cell migration and wound healing, suggesting that the protein might have a role in the regulation of rapid epithelialization during restoration of skin barrier function. These results have important implications for how future therapeutic strategies might be employed to target a multifaceted process such as wound healing. Importantly, a study showed that a non-transcriptional function of Oct4 in the regulation of mitotic entry is crucial for maintaining a short cell cycle in embryonic stem cells, and thus pluripotency^62^. This was achieved by direct inhibition of Cdk1 which resulted in delayed mitotic entry and preservation of embryonic stem cell integrity. We report that absence of Oct4 in skin keratinocytes induced a significantly higher percentage of G2/M-phase cells compared to WT cells, indicating that Oct4 might be implicated in mechanisms that control the G2/M DNA damage checkpoint. Furthermore, we have identified by RNA-seq and ATAC-seq that CDK14, a cyclin-dependent kinase that during mitosis is recruited by CCNY to the plasma membrane and phosphorylates LRP6 receptor of the canonical Wnt pathway^63^, was among the top upregulated genes in Oct4-deficient cells. DIPs of CDK14 demonstrate that Oct4 controls this gene by regulating chromatin regions and possibly transcription factor binding, thus linking Oct4 with cell cycle events.

Tissue morphogenesis and remodeling require cell shape changes and movements that are generated by actin filament organization and actomyosin contractility^26^. Moreover, terminal differentiation of epidermal keratinocytes is associated with actin microfilament reorganization^64^. We present striking evidence that Oct4 is a master organizer of cortical actin in skin keratinocytes with consequence the preservation of epithelial identity. We observed that *OCT4* knockout cells appeared with a plasma membrane where actin cortex was almost absent, increased puncta in the cytoplasm and with the formation of stress fibers. These cells lost their cell-cell contacts and transitioned into a mesenchymal-like phenotype with enlarged size and disorganized actin cytoskeleton. We conclude that loss of Oct4 in skin keratinocytes leads to changes in cellular morphology by triggering cell redistribution of filamentous actin and altering cytoskeletal dynamics. ECM is a complicated network of proteins, fundamental to all organs, with a broad spectrum of functions in tissue homeostasis and remodeling. It provides essential mechanical stability for the skin, through communication with epidermal keratinocytes^65^. Fibronectin is one of the dominant structural components of basement membrane, where it creates a meshwork with collagen type IV and laminin. In addition, fibronectin serves as a mesenchymal marker, with increasing levels during epithelial-to-mesenchymal transition^66^. *OCT4* knockout cells exhibited dramatic increase of fibronectin deposition in ECM compared to WT cells, suggesting that Oct4 controls ECM architecture via regulating the levels of fibronectin in skin keratinocytes.

All layers of the epidermis form a protective barrier from their environment through tight mechanical cohesion between cells of the same or different epidermal layer. Intercellular junctions, such as adherens and tight junctions, are responsible for connecting each cell to each other to establish a tight skin barrier^43^. A recent study showed that overexpression of Oct4 influenced the morphology and adhesion of human hair follicle mesenchymal stem cells^67^. We found that loss of Oct4 in skin keratinocytes diminished E-cadherin and β-catenin from cell-cell contacts by altering their localization, and inhibited their colocalization at the plasma membrane, possibly by impairing the interaction between the two proteins. These observations were accompanied by a significant reduction in cell adhesion, and they were consistent for all clonal cell lines examined. We also demonstrate that ZO1, which belongs to the tight junction family of scaffolding proteins, was lost from the cell periphery of Oct4-deficient cells, indicating that Oct4 controls both adherens and tight junctions to preserve keratinocyte phenotype.

The transcriptional role of Oct4 has been well recognized and documented^60^. To identify whether the observed morphological changes of *OCT4* knockout cells are linked to an altered transcriptional profile, we investigated the transcript levels of various genes involved in the regulation of actin cytoskeleton and ECM. We showed that *E-cadherin*, *ZO1*, *WASP*, *N-WASP*, *CDC42*, *Cofilin-1*, *ROCK1*, *LIMK1*, *Vimentin*, and *β-actin* mRNA levels were significantly increased in Oct4-deficient cells compared to WT keratinocytes, whereas *β-catenin* levels were attenuated in all clonal cells. Moreover, ECM genes such as *FN1*, *Tenascin C* and *TSP1* had also significantly increased transcript levels in *OCT4* knockout cells, with *FN1* and *Tenascin C* exhibiting the most remarkable change. Notably, the decreased relative protein expression of ZO1 and E-cadherin in Oct4-deficient cells does not agree with their mRNA levels. This discrepancy might be attributed to post-transcriptional mechanisms that can shape protein abundance independently of transcript levels, suggesting that Oct4 might regulate the expression and function of these genes through different mechanisms. Our qRT-PCR results align well with the RNA-seq findings, which show that in the absence of Oct4 cytoskeletal and ECM balance is drastically disrupted. Our study defined a core of cytoskeletal, ECM and differentiation related genes, including, *ROBO1*, *CDK14*, *DSG1*, *TGFBR3*, *FGD3*, *TBX18* and *P2YR6* that were deregulated at the mRNA and chromatin accessibility level as a result of loss of Oct4 in skin keratinocytes. For example, *ROBO1* one of the top upregulated genes is involved in directing migration of many cell types^68^, limiting branch formation in the mammary gland by antagonizing canonical Wnt signaling and disorganizing myoepithelial cells^69^. *FGD3* which was significantly downregulated in the intersection set, controls cell shape and regulates the actin cytoskeleton by promoting filopodia formation^58^. *LAMB1* and *TNC* major ECM proteins involved in the regulation of stromal-epithelial interactions that are located in the basement membrane^70^, were downregulated and upregulated in Oct4-deficient cells, respectively. Moreover, *RhoA* a member of the Ras-related family of GTPases critical in the regulation of the cytoskeleton through the assembly of actin stress fibers^71^ was found to be associated with all five pathways of interests, suggesting that Oct4 might play an important role in regulating *RhoA* function. Interestingly, the majority of significant DIPs were downregulated in clone 4, highlighting that Oct4 is predominantly a positive regulator of chromatin accessibility. In this context, a change in the levels of Oct4 defines how cytoskeletal and ECM genes are distributed and function in skin keratinocytes, whereas Oct4 might exert its action by regulating transcription factor binding to chromatin.

Our mechanistic investigation using the transcriptionally active (pCEP-OCT4-AA) and inactive (pCEP-OCT4-EE) Oct4 mutant constructs provided evidence that Oct4 can also maintain keratinocyte phenotype through non-transcriptional functions. Ectopic expression of WT and mutant Oct4 plasmids in clonal cell lines restored keratinocyte characteristics. Furthermore, transfection of *OCT4* knockout cells with pCEP-OCT4-WT or pCEP-OCT4-AA reduced mRNA levels of *WASP*, *CDC42* and *ROCK1* compared to untransfected cells, whereas transfection with the transcriptionally inactive mutant pCEP-OCT4-EE showed no decrease in transcript levels of these genes. These findings indicate that Oct4 might shape the cytoskeletal landscape both through its pioneer transcriptional role but also through non-transcriptional mechanisms that might involve complex protein networking.

Our findings highlight that Oct4 has an unexpected functional role in skin keratinocytes by preserving epithelial identity through dynamic regulation of actin cytoskeleton and by reshaping the transcriptional and chromatin landscapes. This work corroborates previous reports of Oct4 crosstalk with the cytoskeleton, and provides evidence that these are not limited to embryonic stem cells.

## Supporting information

Key Resources Table

## ACKNOWLEDGMENTS

We are grateful to Professor Paris Skourides and Mr. Adonis Hadjigeorgiou for providing assistance with the live imaging microscopy set-up. This work was co-funded by the European Regional Development Fund and the Republic of Cyprus through the Research & Innovation Foundation (projects OPPORTUNITY/0916/ERC-StG/003 and INFRASTRUCTURES/1216/0034), awarded to KS. Graphical abstract was created with BioRender.com.

## AUTHOR CONTRIBUTIONS

E.D.C. and K.S. conceived the study, generated hypotheses and edited the manuscript. E.D.C. drafted the manuscript, assembled the figures, analyzed the data, designed and performed most of the experiments. M.T. performed all the bioinformatics and computational analysis of RNA-seq and ATAC-seq data. G.M.S. provided scientific advice about bioinformatics analysis. C.V. performed computational analysis for the quantification of fluorescence from images obtained during confocal microscopy. C.M. assisted with confocal microscopy. T.S. provided scientific advice and feedback. K.S. supervised the project and acquired funding.

## STAR METHODS

### RESOURCE AVAILABILITY

#### Lead contact

Further information and requests for resources and reagents should be directed to and will be fulfilled by the lead contact, Katerina Strati

#### Materials availability

DNA constructs/plasmids generated by the authors will be distributed upon request to other researchers.

#### Data and code availability

- All datasets used are summarized in the key resources table. Next generation sequencing data generated for this study have been deposited in the National Center for Biotechnology Information (NCBI) gene expression omnibus (GEO) and in sequence read archive (SRA) with accession numbers GSE230655 and PRJNA961975, respectively. SubSeries accession numbers for individual RNA-seq and ATAC-seq datasets are GSE230653 and GSE230654, respectively. Original Western blot images have been deposited at Zenodo and are publicly available as of the date of publication. The DOI is listed in the key resources table.
- R scripts used for filtering, processing and visualization of data are available upon request.
- Any additional information required to reanalyze the data reported in this paper is available from the lead contact upon request.

### EXPERIMENTAL MODEL AND SUBJECT DETAILS

#### Cell lines and culture conditions

HaCaT immortalized keratinocyte cell line from male adult human skin (CLS, 300493) and HeLa human cervical cancer cell line (CLS, 300194) were cultured at 37 °C in 5% CO2, in DMEM (Gibco, 41965039) supplemented with 10% FBS (Gibco, 10500064) and 1% penicillin-streptomycin (Gibco, 15070063). C33a human cervical cancer cell line (ATCC, CRM-HTB-31) was cultured at 37 °C in 5% CO2, in MEM (Gibco, 11090081) supplemented with 10% FBS (Gibco, 10500064), 1% penicillin/streptomycin (Gibco, 15070063) and 1% L-glutamine (Gibco, 25030024). Detachment of HaCaT cells was performed in 0.05% Trypsin-EDTA (Gibco, 25300062), whilst time of detachment is defined as the time when all cells detached from the flask. All cell lines were routinely tested for the absence of mycoplasma.

### METHOD DETAILS

#### Western Blot

Cells were washed twice with ice-cold 1X phosphate buffered saline (PBS) and homogenized in sample buffer (50 mM Tris-HCl; pH 6.8, 2% SDS, 0.1% bromophenol blue, 10% glycerol) supplemented with 200 mM Dithiothreitol (ThermoFisher, R0861) and protease inhibitor tablets (ThermoFisher, A32955) 48 h after seeding. Then, cells were lysed by gentle sonication and heated at 95 °C for 7 min. Protein samples were separated by SDS-PAGE and transferred onto nitrocellulose membranes (Bio-Rad). Mono/polyclonal primary antibodies and appropriate fluorescent-conjugated secondary antibodies were used for blotting. All membranes were incubated with primary antibodies overnight at 4 °C. Protein bands were visualized using the ChemiDoc MP imaging system (Bio-Rad). The following antibodies were used: Oct4 (Cell Signaling, 2750) 1:1000, GAPDH (abcam, 9484) 1:2000, E-cadherin (ThermoFisher, PA5-32178) 1:5000, β-catenin (ThermoFisher, 13-8400) 1:1000, ZO1 (proteintech, 66452-1-Ig) 1:3000, β-actin (abcam, 8226) 1:1000, goat anti-rabbit DyLight-680 (ThermoFisher, 35568) 1:10000, goat anti-mouse DyLight-800 4X PEG (ThermoFisher, SA5-35521) 1:10000, donkey anti-goat DyLight-680 (ThermoFisher, SA5-10090) 1:10000. GAPDH was used as loading control. Protein bands were quantified using ImageJ. Complete blots are available at Zenodo (DOI: 10.5281/zenodo.10223953).

#### Transfection

1 x 10^6^ HaCaT cells were seeded for 24 h. Then, cells were transfected with 5 μg of plasmid pcDNA3-EGFP (addgene, 13031), or pcDNA3.3-OCT4 (addgene, 26816), or pcDNA3-OCT4EGFP, or pCEP4-OCT4-WT (addgene, 40629), or pCEP4-OCT4-T234A-S235A (addgene, 40630) or pCEP4-OCT4-T234E-S235E (addgene, 40631) for 48 h using FuGENE 6 (Promega, E2691) according to the manufacturer’s protocol.

#### Immunoprecipitation

For immunoprecipitation, 6 x10^6^ cells were seeded for 48 h. Untreated or transfected HaCaT cells were washed twice on ice with ice-cold 1X PBS, lysed in RIPA buffer (NaCl 150 mM, EDTA pH=8; 5 Mm, Tris-HCl pH=8; 50 mM, 1% Triton X-100, 0.5% sodium deoxycholate, 0.1% SDS) supplemented with protease inhibitor tablets (ThermoFisher, A32955) for 15 min on ice and centrifuged at 4 °C for 15 min at 12.500 rpm. Supernatants were transferred to a clean tube. Protein G sepharose 4 fast flow beads (Cytiva, 17061801) were washed three times with RIPA buffer and a 50% slurry was prepared. Cell lysates were cleared with 80 μl protein G sepharose beads for 1 h at 4 °C on a rotator. Supernatants were collected after centrifugation at 12.000 rpm at 4 °C. Following clearing step, 10% of the lysate was used as input control. The rest of the lysate was split in half and either 1:100 Oct4 antibody (Cell Signaling, 2750) or rabbit IgG isotype control (Invitrogen, 10500C) was added. Mixtures were incubated overnight at 4 °C on a rotator. Next day, 90 μl of protein G sepharose beads were added in all mixtures, followed by incubation at 4 °C on a rotator for 3 h. Beads were washed three times with ice-cold RIPA buffer and proteins were eluted in sample buffer by boiling at 95 °C for 10 min followed by western blotting analysis.

#### Quantitative real time-PCR (qRT-PCR)

Total RNA was isolated from the cells using RNeasy Maxi Kit (QIAGEN, 75162) as per the manufacturer’s instructions. All RNA samples were treated on column with DNase I (QIAGEN, 79254). First strand cDNA was transcribed (200 ng) with iScript cDNA synthesis kit (Bio-Rad, 1708891). qRT-PCR was performed using a CFX96 Touch Real-Time PCR Detection System Thermal Cycler (Bio-Rad) and KAPA SYBR FAST qPCR Master Mix (2X) kit (KAPA BIOSYSTEMS, KK4602) with first strand cDNA, forward and reverse primers (IDT). The list of primers is given in Table S2. The PCR cycling parameters used were as follows, enzyme activation (95 °C for 2 min), denaturation/annealing (95 °C for 2 sec followed by 60 °C for 20 sec, x40 cycles) and extension (72 °C for 5 sec). Ct values were normalized to endogenous *GAPDH* (housekeeping gene) using the ^ΔΔ^Ct method.

#### Cell size quantification

Cells were seeded in 6-well plates for 48 h (untreated or transfected). Then, images were acquired using phase-contrast microscopy and a 20x objective (Zeiss Observer A1 microscope). Images were analyzed completely blinded. Cell area of single cells (cell by cell quantification) was measured by selecting the cell perimeter using ImageJ software (National Institutes of Health). Minimum of 545 cells were measured per condition.

#### Flow cytometry

For cell size analysis: 2 x 10^6^ HaCaT WT or *OCT4* knockout cells were dissociated using 0.05% trypsin-EDTA (Gibco, 25300062) and washed twice in ice-cold 1X PBS. Cells were passed through a 0.2 μm filter and analyzed using an S3e Cell Sorter (Bio-Rad). Data were analyzed using FlowJo version 10, and gating strategy is shown in Figure S2C.

For cell cycle analysis: 1 x 10^6^ HaCaT WT or *OCT4* knockout cells were dissociated using 0.05% trypsin-EDTA (Gibco, 25300062). Cell pellet was washed twice in 1 ml 1X PBS at room temperature (20-24 °C). Then, 4 mL of absolute ethanol at -20 °C was added dropwise and slowly in the cell suspension while vortexing at top speed. Cells were left in ethanol at -20 °C for 2 h. After ethanol incubation, cells were pelleted by centrifugation at 3500 rpm, and 5 ml of 1X PBS were added allowing cells to rehydrate for 15 min. Then, 1 ml 1X PBS containing 1:500 propidium iodine (PI) (ThermoFisher, P3566) and 1:100 RNase A (MACHEREY-NAGEL, 740505) was added in each sample for 15 min at room temperature. Cells were passed from a 0.2 μm filter and analyzed for cell cycle distribution by flow cytometry in the presence of the dye. Data were analyzed using FlowJo version 10.

#### Cell proliferation

Cells were seeded at a density of 8 x 10^4^ cells per well in 24-well plates, in triplicates, and incubated at 37 °C. Proliferation rates were determined at 24, 48 and 72 h after plating. Quantification of cell number per ml was measured on a TC20 Cell Counter (Bio-Rad).

#### Wound healing assay

For end point cell migration: 1.5 x 10^6^ cells were seeded in 24-well plates in complete DMEM. After 24 h, cell monolayer was washed twice with 1X PBS and scratched by a sterile 200 μl pipette tip. 500 μl of DMEM containing 1% FBS were added in each well. Cells were incubated at 37 °C in 5% CO2 for 20 h. Images were taken at time point zero and after 20 h under an Axiovert 200 motorized inverted microscope (Carl Zeiss AG, Germany) using an Axiocam system and a Zeiss Plan-Apochromat 2.5x objective. Cell migration distances into the scratched area were measured with AxioVision Rel. 4.8 software in 10 randomly chosen fields.

For live cell migration: 2.5 x 10^6^ cells were seeded in a two-well μ-slide/chamber (ibidi, 80286) in complete DMEM, incubated at 37 °C in 5% CO2 for 24 h. After 24 h, cell monolayer was washed twice with 1X PBS and scratched by a sterile 200 μl pipette tip. 1.5 ml Leibovitz’s L-15 medium/CO2 independent (Gibco, 11415064) was added in each well and the chambers were transferred on a stage with a heating system (ibidi, 10927) connected to the microscope mentioned above. The duration of time-lapse image acquisition was 20 h with a capturing interval of 2 h. All images and videos were acquired at 5x magnification (Zeiss Plan-Apochromat 5x objective). For statistical analysis, 5-10 random fields per image were chosen. Gap closure was measured at the exact same field at time point zero and at 20 h using AxioVision Rel. 4.8 software. Movies (AVI format) were prepared using ImageJ software (US National Institutes of Health (NIH)) and converted into MP4 format using Clipchamp video editor (Microsoft).

For the presentation of graph in Figure 2B end-point and time-lapse microscopy data were combined and analyzed together. Gap closure was measured at time point zero and after 20 h.

#### Immunofluorescence microscopy and imaging

HaCaT WT or *OCT4* knockout cells were fixed using 4% paraformaldehyde (PFA) in 1X PBS for 10 min at room temperature. Cells were washed thrice with 1X PBS followed by permeabilization and blocking using 0.1% Triton X-100 (Sigma, T8787) and 2% Bovine Serum Albumin (BSA) (A7906, Sigma) in PBS for 15 min. Cells were then incubated in primary antibodies diluted in 1X PBS containing 2% BSA and 0.1% Triton X-100 for 1 hour at room temperature. The primary antibodies and dye used were: E-cadherin (ThermoFisher, PA5-32178) 1:500, β-catenin (ThermoFisher, 13-8400) 1:300, ZO1 (ThermoFisher, 33-9100) 1:100, fibronectin (Sigma, F3648) 1:500, GFP (SICGEN, AB0020-500) 1:500 and Alexa Fluor Plus 555 conjugated phalloidin (Invitrogen, A30106) 1:400. After washing with 1X PBS thrice, cells were then incubated with secondary antibodies in 1X PBS containing 2% BSA and 0.1% Triton X-100 for 45 min at room temperature. All secondary antibodies were used at 1:400. The secondary antibodies used were: Alexa Fluor 594-conjugated goat anti-mouse IgG (Jackson ImmunoResearch, 115-585-003), FITC goat anti-rabbit IgG (Jackson ImmunoResearch, 111-095-003) and Alexa Fluor 488-conjugated donkey anti-goat IgG (Jackson ImmunoResearch, 705-545-003). Cells were then washed thrice with 1X PBS and mounted using Vectashield with DAPI (VECTOR laboratories, H-1500). Cells were imaged either with a Zeiss Observer A1 microscope and the AxioVision software 4.8 or with a STELLARIS 5 confocal microscope (Leica, Germany) at 40x (HC PL APO 40x/1.30 OIL CS2) or 63x (HC PL APO 63X/1.40 OIL CS2) magnification objectives, using the Leica Application Suite X software.

#### Fluorescence quantification

Fluorescence intensity was analyzed using both custom developed tools and built-in algorithms in MATLAB. Specifically, the mean fluorescence intensity of phalloidin, ZO1, E-cadherin, fibronectin and β-catenin immunostaining was quantified to estimate their overall concentration. This was achieved by identifying regions of interest that correspond to stained cellular structures and by calculating the total number of detected positive pixels based on a predetermined intensity threshold. For β-catenin/E-cadherin colocalization calculations were performed in pixel regions where both colors overlapped. Additionally, fluorescence intensity of phalloidin, ZO1, E-cadherin and β-catenin was measured specifically either on the cell membrane or intracellularly in single cells. The cellular membrane was demarcated using morphological image processing techniques and the average intensity distribution was computed both within the membrane and the central region of the cell. To gain more insight into protein distribution and localization the ratio of membrane mean fluorescence to the cytoplasmic mean fluorescence was measured. Intensity profiles were generated by measuring the intensity of phalloidin, ZO1, E-cadherin, and β-catenin along the cell diameter of single cells. For normalized data sets, all values have been normalized against the average intensity of WT samples. Approximately 140-200 cells per staining were analyzed (14-20 cells per image). Images from three biological experiments were acquired (n=3).

#### Plasmid construction

pcDNA3-OCT4EGFP was constructed by amplifying OCT4 from pcDNA3.3-OCT4 plasmid (addgene, 26816) using the pcDNA3.3OCT4-NEB-F: 5’- caagcttggtaccgagctcggatccATGGGCGGTAGGCGTGTAC and pcDNA3.3OCT4-NEB-R: 5’-ccttgctcaccatctcgagcggccgcgcGTTTGAATGCATGGGAGAGCC primers (IDT). 1ng pcDNA3.3-OCT4 was added for the amplification using the Q5 High-Fidelity DNA Polymerase kit (NEB, M0491). pcDNA3-EGFP (addgene, 13031) vector was digested with NotI-HF (NEB, R3189) and BamHI-HF (NEB, R3136) restriction enzymes. The 1.3 Kb OCT4 insert was ligated into the digested pcDNA3-EGFP vector using the NEBuilder HiFi DNA Assembly Master Mix (NEB, E2621) according to the manufacturer’s instructions. Ligation product was used to transform 5-alpha competent *E. coli* (NEB, C2987I) cells. pcDNA3-OCT4EGFP DNA was purified using a plasmid mini kit (QIAGEN, 12123). Correct orientation of the insert was confirmed by sanger sequencing.

#### CRISPR-Cas9 genome editing

Cas9 ribonucleoproteins (RNPs) were prepared fresh. crRNA and tracrRNA-ATTO 550 (IDT, 1075927) were diluted in nuclease-free duplex buffer (IDT, 11-01-03-01), mixed 1:1 and incubated 5 min at 95 °C to generate 1 μM crRNA:tracrRNA-ATTO 550 duplexes. crRNA sequence: 5’-TATTCCTTGGGGCCACACGT-3’, PAM-AGG (IDT). A 20 nucleotide protospacer sequence crRNA (IDT, 1072544) designed to be non-targeting in human genome was used as a negative control.

For the formation of the RNP complex: Cas9 (IDT, 1081058) was diluted in OptiMEM (ThermoFisher, 51985091) media at 1 μM concentration. Then, 1 μM crRNA:tracrRNA-ATTO 550 duplex was mixed with 1 μM Cas9 enzyme in OptiMEM media, in a 96-well plate, for 5 min at room temperature to assemble the RNP complexes. Following RNP formation, 1.3 μl lipofectamine RNAiMAX (ThermoFisher, 13778100) transfection reagent was added for 20 min to generate the transfection complexes. Low passage HaCaT cells (50.000 cells/ml) were added to the transfection complexes (10 nM RNP final concentration) and incubated at 37 °C with 5% CO2 for 48 h.

After 48 h incubation, genomic DNA was extracted by addition of 50 μl QuickExtract DNA solution (Epicentre, QE09050) to the cells, and incubation in a thermal cycler at 65 °C for 10 min, followed by 98 °C for 5 min. Then, genomic DNA was amplified using KAPA HiFi HotStart PCR kit (Kapa Biosystems, KK2501) according to the manufacturer’s guidelines and specific primers flanking the CRISPR cut site (with the CRISPR cut site off center) that generate a 945 kb amplicon. Primers used: OCT4/T7E1-F: GAGACATGATGCTCTTCCTTT and OCT4/T7E1-R: CCACTAGGTTCAGGGATACT. Cycling parameters used: Initial denaturation (95 °C for 5 min), denaturation (98 °C for 20 sec, x32), annealing (64 °C for 15 sec, x32), extension (72 °C for 30 sec, x32) and final extension (72 °C for 2min). Genome editing and mutation detection was initially tested with the T7E1 assay using the genome editing detection kit (IDT, 1075932) according to the manufacturer’s instructions. T7E1 digested products were visualized on 2% agarose gels (Figure S1A).

### HaCaT *OCT4* knockout clonal cell line generation

After genome editing validation by T7E1 assay, cells reverse transfected with RNP complexes as mentioned above were collected and washed with 1X PBS. The edited cell pool was then sorted onto single cells in 96-well plates, containing complete DMEM, by flow cytometry. Individual clones were expanded and genotyped to analyze the type of edit. Genomic DNA from each *OCT4* knockout clone or WT was extracted, amplified as above, and analyzed by sanger sequencing (Macrogen, Europe) followed by inference of CRISPR editing software (ICE) (Synthego, CA, USA) analysis to validate every clonal edit (Figure S1B). *OCT4* knockout clonal cell lines were used up to passage 20.

### Amplicon sequencing

Genome editing was also determined by amplicon sequencing. The presence of insertions or deletions around the Cas9 targeted sequence was used to examine genome editing efficiency (Figure S1C). Briefly, genomic DNA from each *OCT4* knockout clone or WT was extracted using DNeasy blood and tissue kit (QIAGEN, 69504) according to the manufacturer’s instructions. Then, 60 ng genomic DNA was amplified as above using primers that generate a 421 kb product. Primers used: Oct4-AS-F: ATCCCTTGGATGTGCCAGTT and Oct4-AS-R:

ACTCCTTAGAGGGGAGATGCG. PCR products were purified using QIAquick PCR Purification Kit (Qiagen, 28104). DNA library preparations, sequencing reactions, and initial bioinformatics analysis were conducted at AZENTA Inc. (South Plainfield, NJ, USA). DNA Library Preparation, clustering, and sequencing reagents were used throughout the process using NEBNext Ultra DNA Library Prep kit following the manufacturer’s recommendations (Illumina, San Diego, CA, USA). End repaired adapters were ligated after adenylation of the 3’ends followed by enrichment by limited cycle PCR. DNA libraries were validated on the Agilent TapeStation (Agilent Technologies, Palo Alto, CA, USA), were quantified using Qubit 2.0 Fluorometer (Invitrogen, Carlsbad, CA) and multiplexed in equal molar mass. The pooled DNA libraries were loaded on the MiSeq Illumina instrument according to manufacturer’s instructions. The samples were sequenced using a 2 x 250 paired-end configuration. Image analysis and base calling were conducted by the Illumina Control Software on the Illumina instrument.

Data analysis: The raw Illumina reads were checked for adapters and quality via FastQC. The raw Illumina sequence reads were trimmed of their adapters and nucleotides with poor quality using Trimmomatic v. 0.36. Paired sequence reads were then merged to form a single sequence if the forward and reverse reads were able to overlap. The merged reads were aligned to the reference sequence and variant detection was performed using AZENTA proprietary Amplicon-EZ program.

#### RNA-seq and analysis

To identify genes that are regulated by Oct4, total RNA from WT or *OCT4* knockout clones 1 and 4 was extracted as described above, for RNA sequencing. 1 μg of high-quality total RNA per sample was used for sequencing. RNA sequencing was performed by AZENTA Life Sciences (South Plainfield, NJ, USA). RNA samples were quantified using Qubit 4.0 Fluorometer (Life Technologies, Carlsbad, CA, USA) and RNA integrity was checked with RNA Kit on Agilent 5300 Fragment Analyzer (Agilent Technologies, Palo Alto, CA, USA). RNA sequencing libraries were prepared using the NEBNext Ultra II RNA Library Prep Kit for Illumina following manufacturer’s instructions (NEB, Ipswich, MA, USA). Briefly, mRNAs were first enriched with Oligo(dT) beads. Enriched mRNAs were fragmented according to manufacturer’s instruction. First strand and second strand cDNAs were subsequently synthesized. cDNA fragments were end-repaired and adenylated at 3’-ends, and universal adapters were ligated to cDNA fragments, followed by index addition and library enrichment by limited-cycle PCR. Sequencing libraries were validated using NGS Kit on the Agilent 5300 Fragment Analyzer (Agilent Technologies, Palo Alto, CA, USA) and quantified by using Qubit 4.0 Fluorometer (Invitrogen, Carlsbad, CA). The sequencing libraries were multiplexed and loaded on the flow cell on the Illumina NovaSeq 6000 instrument according to manufacturer’s instructions. The samples were sequenced using a 2 x 150 Pair-End configuration v1.5. Image analysis and base calling were conducted by the NovaSeq Control Software v1.7 on the NovaSeq instrument. Raw sequence data (.bcl files) generated from Illumina NovaSeq was converted into fastq files and de-multiplexed using Illumina bcl2fastq program version 2.20. One mismatch was allowed for index sequence identification. After investigating the quality of the raw data, sequence reads were trimmed to remove possible adapter sequences and nucleotides with poor quality using Trimmomatic v.0.36. The trimmed reads were mapped to the reference genome using the Bowtie2 aligner v.2.2.6. BAM files were generated as a result of this step. Unique gene hit counts were calculated using featureCounts from the Subread package v.1.5.2. Only unique reads that fell within gene regions were counted.

The counts from all samples were used to contract the raw counts table which was used for downstream differential expression analysis. Using DESeq2, a comparison of gene expression between WT vs clone 1 and WT vs clone 4 was performed. The Wald test was used to generate p-values and log2 fold changes. Genes with an adjusted p-value < 0.05 and absolute log2 fold change > 1 were called as differentially expressed genes for each comparison.

#### ATAC-seq and analysis

For ATAC sequencing analysis, 1 x10^6^ cells from WT or *OCT4* knockout clone 4 were washed with 1X PBS and pelleted. Cell viability was evaluated using LIVE/DEAD fixable dead cell stain kit (ThermoFisher, L34969) with flow cytometry. Cells with <90% viability were used for ATAC sequencing. ATAC-seq library preparations, sequencing reactions and initial bioinformatic analysis were conducted at AZENTA Life Sciences (South Plainfield, NJ, USA) as follows: Live cell samples were thawed, washed, and treated with DNAse I (LifeTech, EN0521) to remove genomic DNA contamination. Live cell samples were quantified and assessed for viability using a Countess Automated Cell Counter (ThermoFisher, Waltham, MA, USA). After cell lysis and cytosol removal, nuclei were treated with Tn5 enzyme (Illumina, 20034197) for 30 minutes at 37°C and purified with Minelute PCR Purification Kit (Qiagen, 28004) to produce tagmented DNA samples. Tagmented DNA was barcoded with Nextera Index Kit v2 (Illumina, FC-131-2001) and amplified via PCR prior to a SPRI Bead cleanup to yield purified DNA libraries. The sequencing libraries were clustered on a single lane of a flowcell. After clustering, the flowcell was loaded on the Illumina HiSeq instrument 4000 according to manufacturer’s instructions. The samples were sequenced using a 2 x 150bp Paired End configuration. Image analysis and base calling were conducted by the HiSeq Control Software (HCS). Raw sequence data (.bcl files) generated from Illumina HiSeq was converted into fastq files and de-multiplexed using Illumina’s bcl2fastq 2.17 software. One mismatch was allowed for index sequence identification.

After investigating the quality of the raw data, sequencing adapters and low-quality bases were trimmed using Trimmomatic 0.38. Cleaned reads were then aligned to reference genome hg38 using bowtie2. Aligned reads were filtered using samtools 1.9 to keep alignments that have a minimum mapping quality of 30, were aligned concordantly and were the primary called alignments. PCR or optical duplicates were marked using Picard 2.18.26 and removed. Prior to peak calling, reads mapping to mitochondria were called and filtered, and reads mapping to unplaced contigs were removed.

MACS2 2.1.2 was used for peak calling to identify open chromatin regions. Valid peaks from each condition were then merged and peaks called in at least 66% of samples were kept for downstream analyses. Reads falling beneath peaks were counted in all samples, and these counts were used for differential peak analyses using the R package Diffbind.

#### RNA-seq and ATAC-seq post-processing analysis

All downstream analysis of the differential expression results was carried out using custom written scripts in R programming language V.4.2.1 and third-party packages. All visualizations were generated using the ggplot2 package and addons in R. Specifically, the volcano plots were generated using the EnhancedVolcano package, using the log2(Fold Change) and -Log10(adjusted p-value) as calculated by DeSeq2 differential expression analysis. The intersection set was generated by finding the common differentially expressed genes between the two RNA-seq comparisons and the genes that also exhibited difference in peaks (FDR<0.05) that overlap either their open reading frames or promoter regions. The ggvenn package was used to plot venn diagrams for the DEGs and DIPs sets and corresponding pathway sets. The sample to gene heatmap was generated using the pheatmap package following a Variance Stabilization Transformation (VST) of the DeSeq2 RNA expression object. Then, the significant DEGs of the intersection set were ranked with respect to their mean abundance across all samples (RNA expressions for WT, clone 1 and clone 4) and the top 10% of these genes were selected to perform the hierarchical clustering analysis. The row- and column-wise hierarchical clustering was performed using the Euclidean distance as a metric and ward.D2 clustering method. Similarly, the sample-to-sample heatmap was generated using the Euclidean pairwise distances calculated on the full count vectors of the samples.

The ATAC-seq peaks were visualized with the IGV web application (www.igv.org/app) on hg38 reference genome using as an input the WT and Clone 4 bigWig track files, as well as the bed file with the regions of significant peaks (FDR<0.01). The number of peaks overlapping the open reading frame and the 3kb upstream of the TSS for selected genes of interest were aggregated and visualized as a barplot using ggplot2. The same package was used to generate the ATAC-seq localization pie chart based on the annotations obtained for the DIPs (FDR<0.05).

All pathway enrichments were calculated using the Overrepresentation Analysis (ORA) method implemented in the clusterProfiler package in R. As an input we used the intersection gene set against the KEGG pathways database. For the latter, genes were mapped from their ENSEMBL to their ENTREZ identifiers. In addition, ORA enrichments were performed using the gene sets from individual comparisons to assess the common and unique pathways enriched from each set. The network of the pathways of interest was generated using the igraph package in R. For the network representation, nodes correspond to pathways of interest and their gene members, while the edges denote the pathway membership.

#### Protein Atlas data re-analysis

Tissue level expression analysis was based on the publicly available data from Protein Atlas (https://www.proteinatlas.org/about/download). The normalized Transcript per Million (nTPM) value was plotted against each tissue using ggplot2. Similarly, we produced the barplots from single cell RNA-seq datasets. The nTPM values for each cell type were obtained again from Protein Atlas (https://www.proteinatlas.org/download/rna_tissue_consensus.tsv.zip, https://www.proteinatlas.org/download/rna_single_cell_type.tsv.zip).

### QUANTIFICATION AND STATISTICAL ANALYSIS

All experiments were repeated at least three times as indicated in the Figure legends. For RNA-seq and ATAC-seq experiments duplicate samples for each condition were analyzed. Images are representative of at least three independent experiments. Statistical analysis and p value determinations were carried out using GraphPad Prism 8 (GraphPad, San Diego, CA). Statistical significance was analyzed by unpaired two-tailed Student’s t-test or one-way analysis of variance (ANOVA) as specified in the respective Figure legends. Data are presented as the mean ± SEM. The significance threshold for all statistical calculations was set as p<0.05 (*p<0.05, **p<0.01, ***p<0.001, ****p<0.0001, ns=non significant, n=number of biological replicates).

## SUPPLEMENTAL ITEM TITLES AND LEGENDS

**Movie S1. Live cell migration of WT skin keratinocytes. Related to Figure 2.**

Representative field of WT keratinocytes showing time-lapse migration. Scale bar, 100 μm.

Movie S2. Live cell migration of *OCT4* knockout clone 1 cell line. Related to Figure 2.

Representative field of clone 1 showing time-lapse migration. Scale bar, 100 μm.

Movie S3. Live cell migration of *OCT4* knockout clone 3 cell line. Related to Figure 2.

Representative field of clone 3 showing time-lapse migration. Scale bar, 100 μm.

Movie S4. Live cell migration of *OCT4* knockout clone 4 cell line. Related to Figure 2.

Representative field of clone 4 showing time-lapse migration. Scale bar, 100 μm.

