## Supplementary material for "Oct4 is a gatekeeper of epithelial identity by regulating cytoskeletal organization in skin keratinocytes": Key Resources Table

| REAGENT or RESOURCE | SOURCE | IDENTIFIER |
|---|---|---|
| **Antibodies** | | |
| Anti-rabbit Oct4 Polyclonal | Cell Signaling Technology | Cat# 2750; RRID:AB_823583 |
| Anti-mouse GAPDH Monoclonal | Abcam | Cat# ab9484; RRID:AB_307274 |
| Anti-mouse β-actin Monoclonal | Abcam | Cat# ab8226; RRID:AB_306371 |
| Anti-rabbit E-cadherin Polyclonal | Thermo Fisher Scientific | Cat# PA5-32178; RRID:AB_2549651 |
| Anti-mouse β-catenin Monoclonal, clone: CAT-5H10 | Thermo Fisher Scientific | Cat# 13-8400; RRID:AB_2533039 |
| Anti-mouse ZO1 Monoclonal, clone: ZO1-1A12 | Thermo Fisher Scientific | Cat# 33-9100; RRID:AB_2533147 |
| Anti-rabbit Fibronectin | Sigma | Cat# F3648; RRID:AB_476976 |
| Anti-goat GFP | SICGEN | Cat# AB0020; RRID:AB_2333100 |
| Anti-mouse ZO1 Monoclonal, clone: 1G4A1 | proteintech | Cat# 66452-1-Ig; RRID:AB_2881821 |
| Goat Anti-Mouse IgG (H+L) Alexa Fluor 594-AffiniPure Secondary Antibody | Jackson ImmunoResearch Labs | Cat# 115-585-003; RRID:AB_2338871 |
| Goat Anti-Rabbit IgG (H+L) Fluorescein (FITC)-AffiniPure Secondary Antibody | Jackson ImmunoResearch Labs | Cat# 111-095-003; RRID:AB_2337972 |
| Donkey Anti-Goat IgG (H+L) Alexa Fluor 488-AffiniPure Secondary Antibody | Jackson ImmunoResearch Labs | Cat# 705-545-003; RRID:AB_2340428 |
| Goat anti-Rabbit IgG (H+L) Secondary Antibody, DyLight™ 680 | Thermo Fisher Scientific | Cat# 35568; RRID:AB_614946 |
| Goat anti-Mouse IgG (H+L) Secondary Antibody, DyLight™ 800 4X PEG | Thermo Fisher Scientific | Cat# SA5-35521; RRID:AB_2556774 |
| Rabbit IgG Isotype Control | Thermo Fisher Scientific | Cat# 10500C; RRID:AB_2532981 |
| Donkey anti-Goat IgG (H+L) Secondary Antibody, DyLight™ 680 | Thermo Fisher Scientific | Cat# SA5-10090; RRID AB_2556670 |
| **Bacterial and virus strains** | | |
| 5-alpha chemically competent *E. coli* cells | New England BioLabs | Cat# C2987I |
| **Chemicals, peptides, and recombinant proteins** | | |
| DMEM | Gibco | Cat# 41965039 |
| MEM | Gibco | Cat# 11090081 |
| FBS | Gibco | Cat# 10500064 |
| 0.05% trypsin-EDTA | Gibco | Cat# 25300062 |
| Penicillin/Streptomycin | Gibco | Cat# 15070063 |
| L-glutamine | Gibco | Cat# 25030024 |
| Leibovitz's L-15 medium/CO2 independent | Gibco | Cat# 11415064 |
| Dithiothreitol | Thermo Fisher Scientific | Cat# R0861 |
| Protease Inhibitor Tablets | Thermo Fisher Scientific | Cat# A32955 |
| FuGENE6 | Promega | Cat# E2691 |
| G Sepharose 4 fast flow beads | Cytiva | Cat# 17061801 |
| DNase I | QIAGEN | Cat# 79254 |
| Propidium Iodine | Thermo Fisher Scientific | Cat# P3566 |

| | | |
|---|---|---|
| RNase A | MACHEREY-NAGEL | Cat# 740505 |
| Triton X-100 | Sigma | Cat# T8787 |
| Bovine Serum Albumin | Sigma | Cat# A7906 |
| Alexa Fluor Plus 555 conjugated phalloidin | Invitrogen | Cat# A30106 |
| Vectashield with DAPI | VECTOR Laboratories | Cat# H-1500 |
| Nuclease-free duplex buffer | IDT | Cat# 11-01-03-01 |
| Cas9 | IDT | Cat# 1081058 |
| OptiMEM | Thermo Fisher Scientific | Cat# 51985091 |
| Lipofectamine RNAiMAX | Thermo Fisher Scientific | Cat# 13778100 |
| QuickExtract DNA solution | Epicentre | Cat# QE09050 |
| **Critical commercial assays** | | |
| iScript cDNA synthesis kit | Bio-Rad | Cat# 1708891 |
| KAPA SYBR FAST qPCR Master Mix (2X) kit | KAPA BIOSYSTEMS | Cat# KK4602 |
| Q5 High-Fidelity DNA Polymerase kit | New England BioLabs | Cat# M0491 |
| NotI-HF | New England BioLabs | Cat# R3189 |
| BamHI-HF | New England BioLabs | Cat# R3136 |
| NEBuilder HiFi DNA Assembly Master Mix | New England BioLabs | Cat# E2621 |
| Plasmid mini kit | QIAGEN | Cat# 12123 |
| KAPA HiFi HotStart PCR kit | KAPA BIOSYSTEMS | Cat# KK2501 |
| Genome Editing Detection kit | IDT | Cat# 1075932 |
| DNeasy blood and tissue kit | QIAGEN | Cat# 69504 |
| QIAquick PCR Purification Kit | QIAGEN | Cat# 28104 |
| LIVE/DEAD fixable dead cell stain kit | Thermo Fisher Scientific | Cat# L34969 |
| **Deposited data** | | |
| RNA-seq and ATAC-seq SuperSeries number | This study | GSE230655 |
| RNA-seq data | This study | GSE230653 |
| ATAC-seq data | This study | GSE230654 |
| Amplicon-seq data | This study | NIH BioProject ID: PRJNA961975 |
| Original Western Blot Images | This study | DOI: 10.5281/zenodo.10223953 (Zenodo) |
| **Experimental models: Cell lines** | | |
| HaCaT cells | CLS | Cat# 300493 |
| HeLa cells | CLS | Cat# 300194 |
| C33a cells | ATCC | Cat# CRM-HTB-31 |
| **Oligonucleotides** | | |
| pcDNA3.3-OCT4-NEB-F: caagcttggtaccgagctcggatccATGGCGGTAGGCGTGTAC | IDT | N/A |
| pcDNA3.3-OCT4-NEB-R: ccttgctcaccatctcgagcggccgcgcGTTTGAATGCATGGGAGAGCC | IDT | N/A |
| Negative control crRNA | IDT | Cat# 1072544 |
| crRNA: TATTCCTTGGGGCCACACGT | IDT | N/A |
| tracrRNA-ATTO 550 | IDT | Cat# 1075927 |
| OCT4/T7E1-F: GAGACATGATGCTCTTCCTTT | IDT | N/A |

| | | |
|---|---|---|
| OCT4/T7E1-R: CCACTAGGTTCAGGGATACT | IDT | N/A |
| Oct4-AS-F: ATCCCTTGGATGTGCCAGTT | IDT | N/A |
| Oct4-AS-R: ACTCCTTAGAGGGGAGATGCG | IDT | N/A |
| Primers for qRT-PCR (see Table S2) | IDT | N/A |
| **Recombinant DNA** | | |
| pcDNA3-EGFP | N/A | RRID: Addgene_13031 |
| pcDNA3.3-OCT4 | Warren et al.[37] | RRID: Addgene_26816 |
| pcDNA3-OCT4EGFP | This study | N/A |
| pCEP4-WT-OCT4 | Brumbaugh et al.[59] | RRID: Addgene_40629 |
| pCEP4-OCT4-T234A-S235A | Brumbaugh et al.[59] | RRID: Addgene_40630 |
| pCEP4-OCT4-T234E-S235E | Brumbaugh et al.[59] | RRID: Addgene_40631 |
| **Software and algorithms** | | |
| Fiji (ImageJ) | NIH, USA | https://fiji.sc/ RRID:SCR_002285 |
| GraphPad Prism 8 | GraphPad Software, Inc | https://www.graphpad.com/ RRID:SCR_002798 |
| AxioVision software 4.8 | Carl Zeiss Microscopy | https://www.micro-shop.zeiss.com/en/us/system/software+axiovision-axiovision+program-axiovision+software/10221/ |
| Inference of CRISPR Edits (ICE) | Synthego | https://ice.synthego.com |
| MATLAB | Mathworks | RRID:SCR_001622 |
| R (v4.2.3) | R core team | https://www.R-project.org/ |
| DESeq2 (v1.38.3) | Love et al.[74] | https://bioconductor.org/packages/release/bioc/vignettes/DESeq2/inst/doc/DESeq2.html |
| Bowtie2 (v2.2.6) | Langmead et al.[75] | https://github.com/BenLangmead/bowtie2 |
| MACS2 (v2.1.2) | Zhang et al.[76] | https://pypi.org/project/MACS2/ |
| IGV webapp (v1.13.3) | Robinson et al.[77] | https://igv.org/app/ |
| DiffBind (v3.8.4) | Bioconductor | https://bioconductor.org/packages/release/bioc/html/DiffBind.html |
| Trimmomatic (v0.38) | Bolger et al.[78] | https://github.com/usadellab/Trimmomatic |
| Samtools (v1.9) | Danecek et al.[79] | https://github.com/samtools/samtools |
| clusterProfiler (v4.6.2) | Wu et al.[80] | https://github.com/YuLab-SMU/clusterProfiler |
