## Supplemental Information for "Oct4 is a gatekeeper of epithelial identity by regulating cytoskeletal organization in skin keratinocytes"

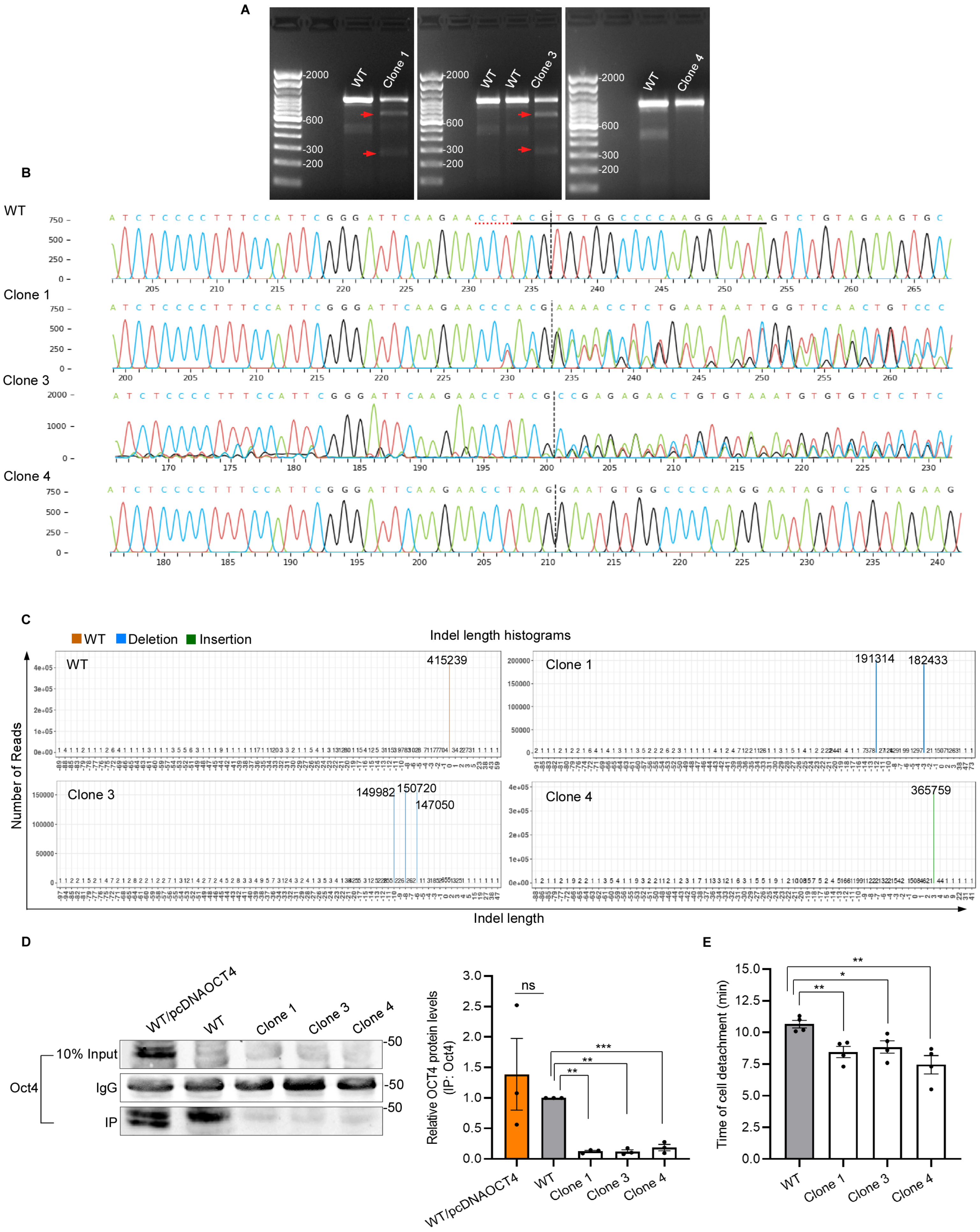

A
WT
Clone 1
Clone 3
Clone 4
2000
600
300
200
B
WT
Clone 1
Clone 3
Clone 4
C
WT
Deletion
Insertion
Indel length histograms
415239
191314
182433
149982
150720
147050
365759
Number of Reads
Indel length
D
WT/pcDNAOCT4
WT
Clone 1
Clone 3
Clone 4
10% Input
IgG
IP
Oct4
-50
Relative OCT4 protein levels (IP: Oct4)
ns
**
**
***
E
Time of cell detachment (min)
**
*
**

**Figure S1. Validation of *OCT4* knockout clones generated by CRISPR-Cas9. Related to Figure 1.**

(A) Representative agarose gel images of T7E1-treated PCR products amplified from the target sites of WT cells (treated with negative control) and edited *OCT4* knockout clones. Red arrows show the generation of specific and predictable size of cleavage products by T7E1. Cleavage does not occur in clone 4 homozygous mutant (right image).

(B) Sanger sequencing chromatograms demonstrating nucleotide alterations after the crRNA cut site in *OCT4* knockout clones compared to WT, due to indel formation. Black line: crRNA sequence, dotted vertical line: cut site, dotted red horizontal line: protospacer adjacent motif. Deconvolution of sanger sequencing traces was performed using Synthego's ICE analysis tool.

(C) Indel quantification and genome editing efficiency was determined by amplicon sequencing. Histograms of WT and *OCT4* knockout clones show number of reads and indel length. Analysis was performed by Genewiz/Azenta.

(D) WT cells transfected with pcDNAOCT4, WT cells (endogenous Oct4) or *OCT4* knockout clones were immunoprecipitated using Oct4 antibody. The immunocomplexes were then eluted and immunoblotted for Oct4 protein. Normal immunoglobulin G (IgG) was used as the negative control and 10% whole lysates (Input) was used as the positive control. Relative Oct4 protein levels were quantified by ImageJ. n=3, unpaired Student's t-test.

(E) WT keratinocytes and *OCT4* knockout clones were detached from the flask at 85-90% confluency using trypsin. Time of cell detachment was measured and set as the time when all cells detached from the flask. n=4, unpaired Student's t-test.

See also Table S1.

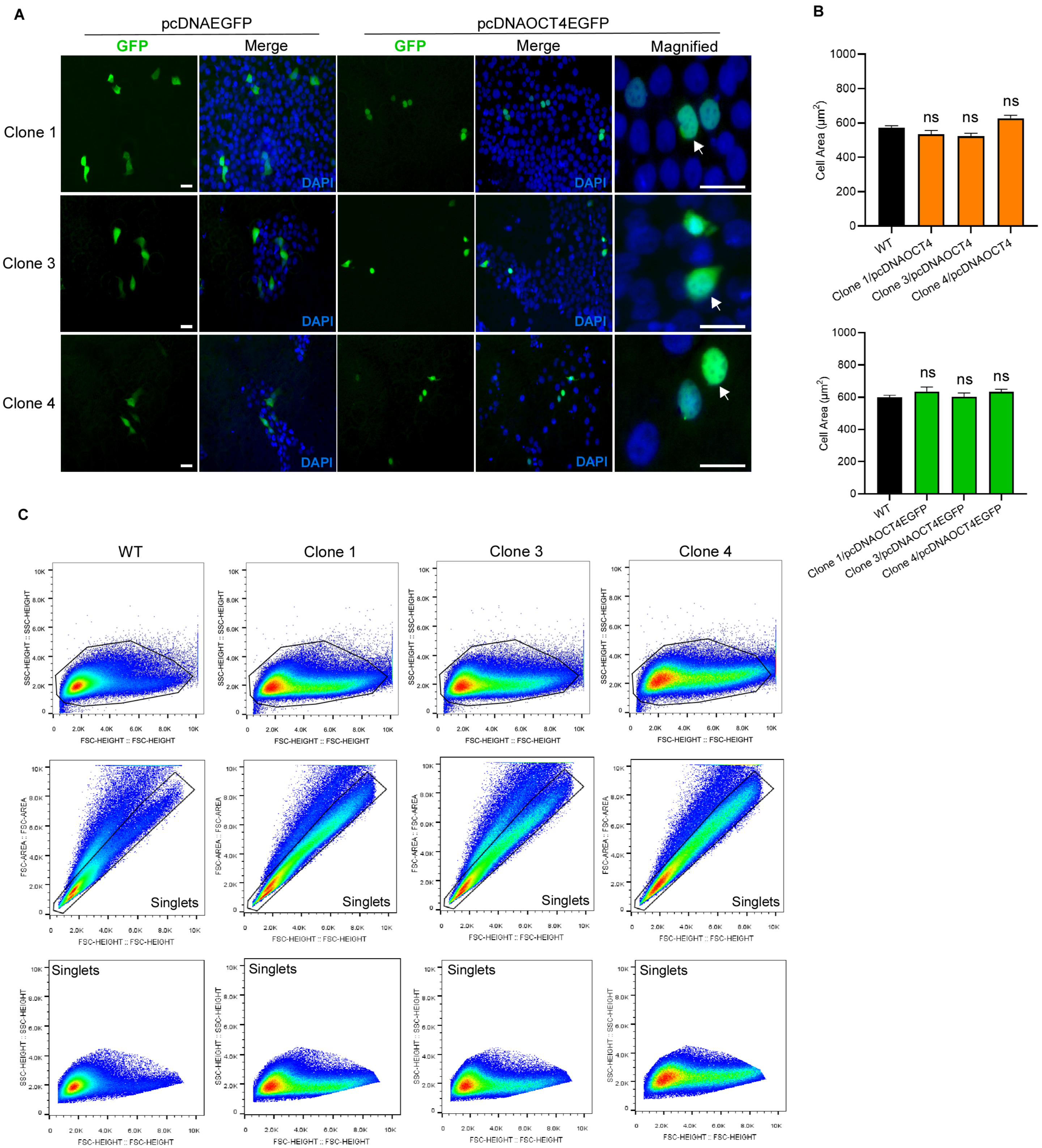
A
pcDNAEGFP
pcDNAOCT4EGFP
GFP
Merge
GFP
Merge
Magnified
Clone 1
Clone 3
Clone 4
DAPI
B
Cell Area (μm²)
1000
800
600
400
200
0
ns
WT
Clone 1/pcDNAOCT4
Clone 3/pcDNAOCT4
Clone 4/pcDNAOCT4
WT
Clone 1/pcDNAOCT4EGFP
Clone 3/pcDNAOCT4EGFP
Clone 4/pcDNAOCT4EGFP
C
WT
Clone 1
Clone 3
Clone 4
SSC-HEIGHT :: SSC-HEIGHT
FSC-HEIGHT :: FSC-HEIGHT
FSC-AREA :: FSC-AREA
Singlets
0
2.0K
4.0K
6.0K
8.0K
10K

**Figure S2. pcDNAOCT4EGFP construct is expressed in all *OCT4* knockout clones and mainly localized in the nucleus. Related to Figure 1.**

(A) Transient expression of pcDNAEGFP (control) or pcDNAOCT4EGFP plasmids in *OCT4* knockout clones 48 h after transfection. Oct4 protein is mainly localized in the nucleus as indicated by white arrows (magnified panel). DAPI was used as nuclear stain. Scale bars, 20 μm.

(B) Cell size measurement of WT keratinocytes compared to *OCT4* knockout clones, transfected either with pcDNAOCT4 (upper) or pcDNAOCT4EGFP (lower) constructs. In clonal cell lines only cells that visually looked smaller and phenotypically altered in the whole cell population were measured. n=3, one-way ANOVA.

(C) Cell size of WT keratinocytes and *OCT4* knockout clones was assessed by flow cytometry. Gating strategy is shown in the forward and side scatter density plots for the generation of the overlay images in Figure 1H.

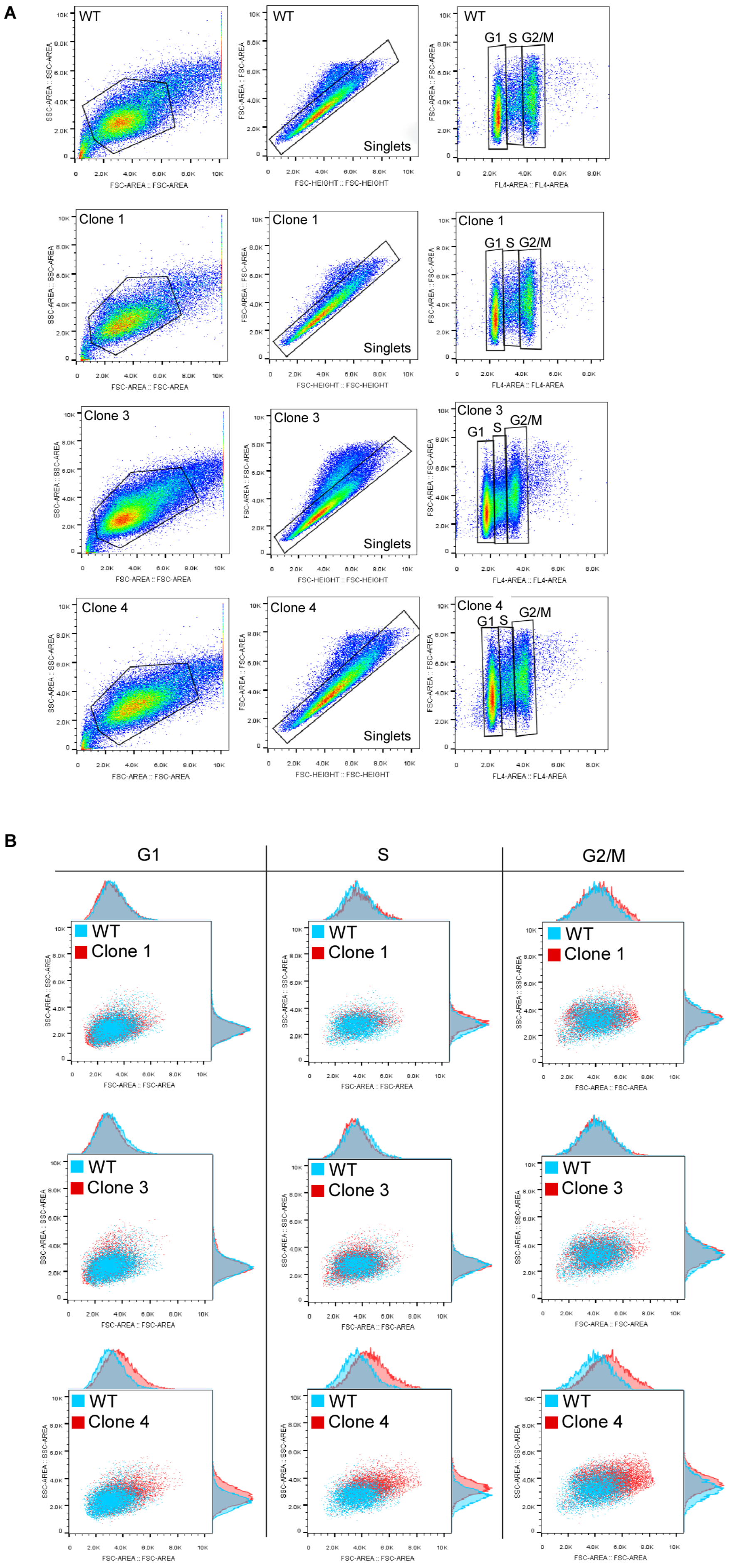
A
WT
Clone 1
Clone 3
Clone 4
Singlets
G1
S
G2/M
SSC-AREA :: SSC-AREA
FSC-AREA :: FSC-AREA
FSC-HEIGHT :: FSC-HEIGHT
FL4-AREA :: FL4-AREA
B
G1
S
G2/M
WT
Clone 1
Clone 3
Clone 4

**Figure S3. Oct4 knockout cells with larger size are equally distributed in cell cycle phases. Related to Figure 2.**

(A) WT and *OCT4* knockout cells stained with PI for cell size analysis in G1, S, and G2/M cell cycle phases. Representative density plots show the gating strategy for the separation of cell cycle phases. n=3.

(B) Cell size comparison between WT keratinocytes and *OCT4* knockout clones in cell cycle phases was assessed by flow cytometry. Overlay images indicate the cell size (forward scatter) and complexity (side scatter) distribution of *OCT4* knockout clones compared to WT keratinocytes in G1, S and G2/M cell cycle phases.

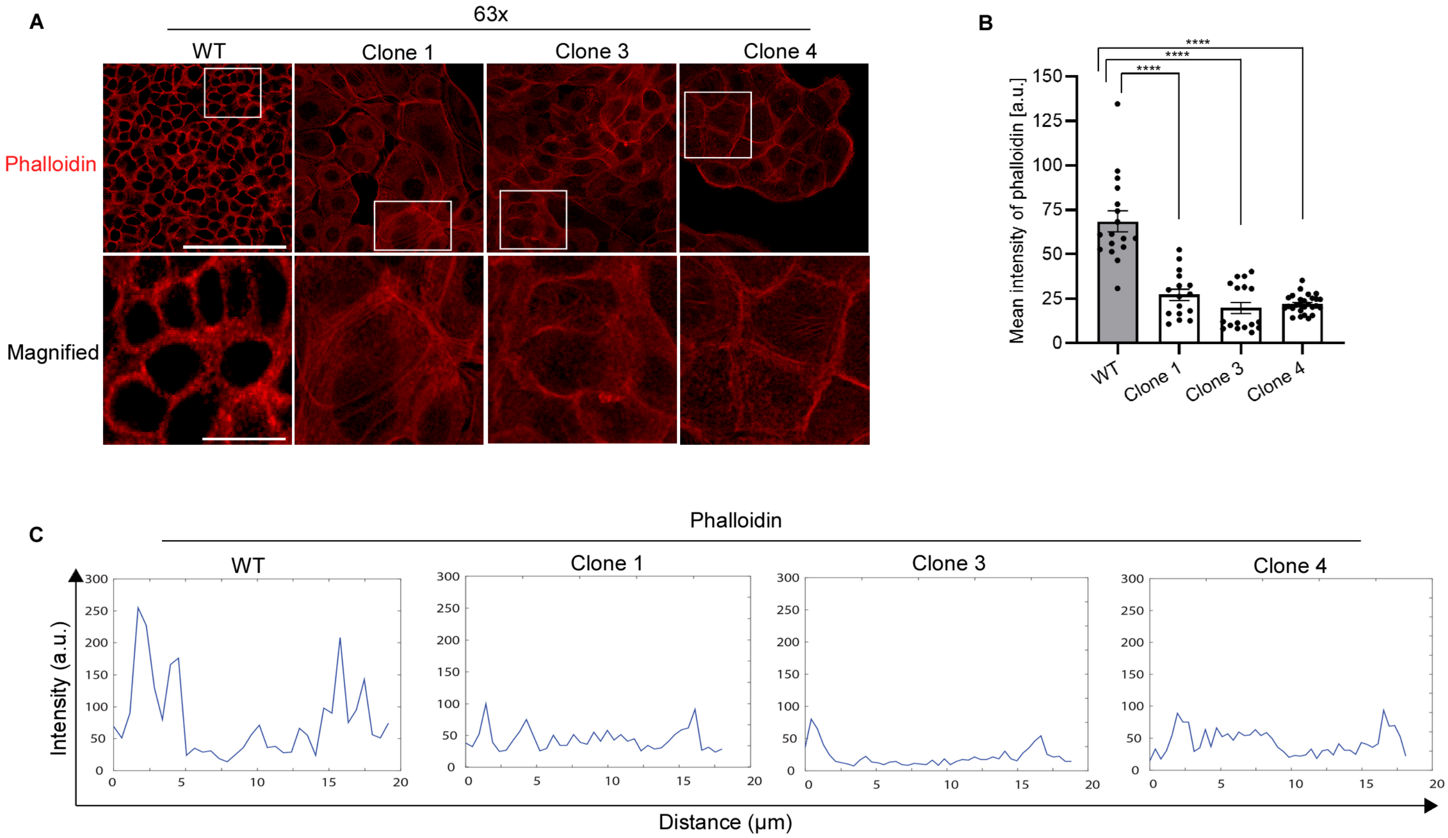
A
63x
WT
Clone 1
Clone 3
Clone 4
Phalloidin
Magnified
B
Mean intensity of phalloidin [a.u.]
****
****
****
0
25
50
75
100
125
150
WT
Clone 1
Clone 3
Clone 4
C
Phalloidin
WT
Clone 1
Clone 3
Clone 4
Intensity (a.u.)
Distance (µm)

**Figure S4. Oct4 knockout affected cortical actin localization and distribution in skin keratinocytes. Related to Figure 3.**

(A) Additional confocal images of WT keratinocytes and *OCT4* knockout clones stained with phalloidin for F-actin visualization at 63x magnification. White boxes indicate magnified image. DAPI is omitted for simplicity. Scale bars, 100 μm and 25 μm for magnified images.

(B) Quantification of phalloidin mean fluorescence intensity comparing WT keratinocytes to *OCT4* knockout clones. n=3, one-way ANOVA.

(C) Representative intensity profiles that were generated by measuring the intensity of phalloidin along the cell diameter. Comparison of fluorescence intensity between WT keratinocytes and *OCT4* knockout clones. These intensity plots provide valuable information on the distribution of F-actin expression within the cell. Specifically, the intensity distribution of F-actin along the diameter of the cell. The x-axis represents the position (membrane-intracellular-membrane) along the cell diameter, while the y-axis represents the intensity of the protein staining. The peaks indicate areas of high protein expression, while the valleys areas of low expression.

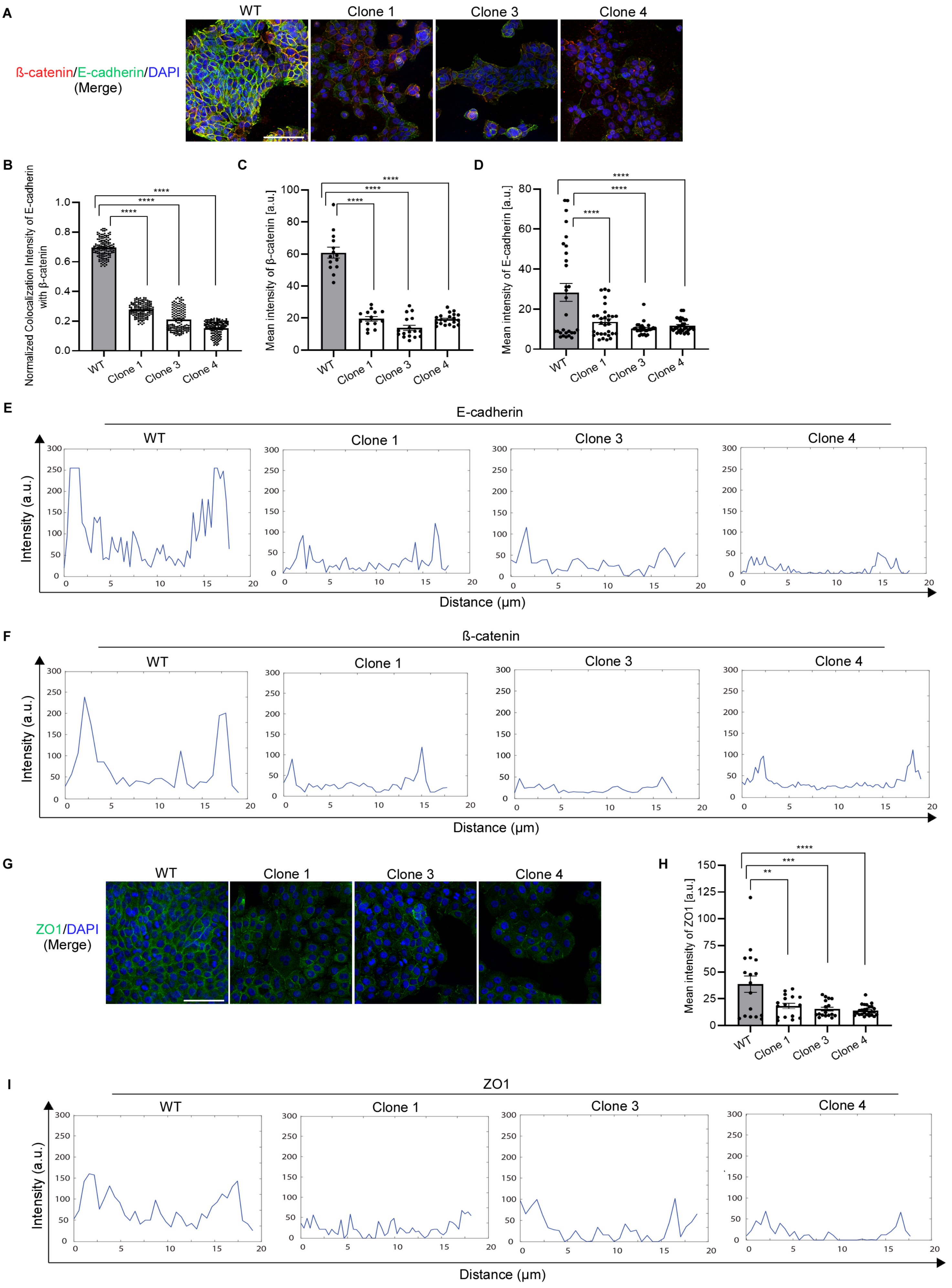

A
WT
Clone 1
Clone 3
Clone 4
ß-catenin/E-cadherin/DAPI
(Merge)
B
Normalized Colocalization Intensity of E-cadherin with β-catenin
1.0
0.8
0.6
0.4
0.2
0.0
****
WT
Clone 1
Clone 3
Clone 4
C
Mean intensity of β-catenin [a.u.]
100
80
60
40
20
0
D
Mean intensity of E-cadherin [a.u.]
E
E-cadherin
Intensity (a.u.)
Distance (μm)
F
ß-catenin
G
ZO1/DAPI
(Merge)
H
Mean intensity of ZO1 [a.u.]
150
125
100
75
50
25
**
***
I
ZO1

**Figure S5. Oct4 regulates adherens and tight junctions in skin keratinocytes. Related to Figure 4.**

(A) Additional confocal images of WT keratinocytes and *OCT4* knockout clones showing colocalization of E-cadherin with β-catenin. Scale bars, 100 μm.

(B) Quantification of β-catenin mean fluorescence intensity comparing WT keratinocytes to *OCT4* knockout clones showing protein concentration. n=3, one-way ANOVA.

(C) Quantification of E-cadherin mean fluorescence intensity comparing WT keratinocytes to *OCT4* knockout clones. n=3, one-way ANOVA.

(D) Quantification of fluorescence intensity of β-catenin with E-cadherin colocalization comparing WT keratinocytes to *OCT4* knockout clones. Colocalization measurements were performed in pixel regions where both colours overlapped. n=3, one-way ANOVA.

(E) Representative intensity profiles showing tracks of a cell that were generated by measuring the intensity of E-cadherin along the cell diameter. Comparison of fluorescence intensity between WT keratinocytes and *OCT4* knockout clones.

(F) Representative intensity profiles that were generated by measuring the intensity of β-catenin along the cell diameter. Comparison of fluorescence intensity between WT keratinocytes and *OCT4* knockout clones.

(G) Additional confocal images of WT keratinocytes and *OCT4* knockout clones indicating ZO1 staining. Merged images with DAPI nuclear stain. Scale bars, 100 μm.

(H) Quantification of ZO1 mean fluorescence intensity comparing WT keratinocytes to *OCT4* knockout clones showing protein concentration. n=3, one-way ANOVA.

(I) Representative intensity profiles showing tracks of a cell that were generated by measuring the intensity of ZO1 along the cell diameter. Comparison of fluorescence intensity between WT keratinocytes and *OCT4* knockout clones.

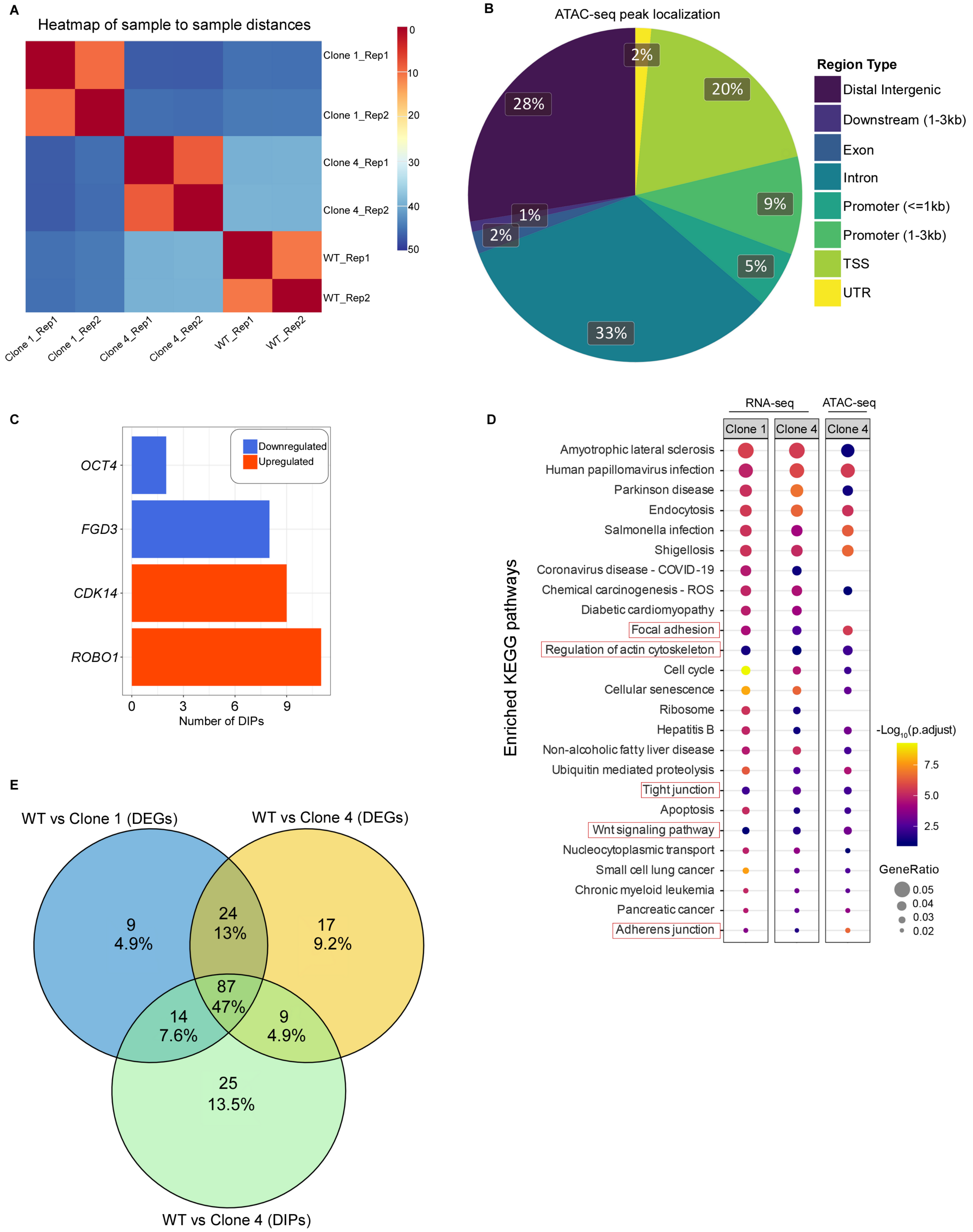
A
Heatmap of sample to sample distances
Clone 1_Rep1
Clone 1_Rep2
Clone 4_Rep1
Clone 4_Rep2
WT_Rep1
WT_Rep2
0
10
20
30
40
50
B
ATAC-seq peak localization
2%
20%
28%
9%
1%
2%
5%
33%
Region Type
Distal Intergenic
Downstream (1-3kb)
Exon
Intron
Promoter (<=1kb)
Promoter (1-3kb)
TSS
UTR
C
OCT4
FGD3
CDK14
ROBO1
Downregulated
Upregulated
0
3
6
9
Number of DIPs
D
RNA-seq
ATAC-seq
Clone 1
Clone 4
Clone 4
Amyotrophic lateral sclerosis
Human papillomavirus infection
Parkinson disease
Endocytosis
Salmonella infection
Shigellosis
Coronavirus disease - COVID-19
Chemical carcinogenesis - ROS
Diabetic cardiomyopathy
Focal adhesion
Regulation of actin cytoskeleton
Cell cycle
Cellular senescence
Ribosome
Hepatitis B
Non-alcoholic fatty liver disease
Ubiquitin mediated proteolysis
Tight junction
Apoptosis
Wnt signaling pathway
Nucleocytoplasmic transport
Small cell lung cancer
Chronic myeloid leukemia
Pancreatic cancer
Adherens junction
Enriched KEGG pathways
-Log10(p.adjust)
7.5
5.0
2.5
GeneRatio
0.05
0.04
0.03
0.02
E
WT vs Clone 1 (DEGs)
WT vs Clone 4 (DEGs)
9
4.9%
24
13%
17
9.2%
87
47%
14
7.6%
9
4.9%
25
13.5%
WT vs Clone 4 (DIPs)

**Figure S6. Loss of Oct4 in skin keratinocytes alters cytoskeletal organization through modulation of chromatin and transcription. Related to Figure 6.**

(A) Heatmap showing the Euclidean distance of the full expression vectors, obtained from the RNA-seq datasets, for WT samples and *OCT4* knockout clone 1 and clone 4 cell lines.

(B) Pie chart of the localization annotations obtained using the DIPs identified in the ATAC-seq dataset (WT compared to clone 4).

(C) Bar plot showing the number of significant DIPs, which fall within the ORF and promoter regions (up to 3 kb upstream of the TSS), of *ROBO1*, *CDK14*, *FGD3* and *OCT4* genes.

(D) ORA enriched KEGG pathways identified using the DEGs from the RNA-seq comparisons (WT compared to clone 1 or clone 4) and the DIPs from the ATAC-seq comparison (WT compared to clone 4). The size corresponds to the gene ratio while the color map indicates the significance of each enriched pathway.

(E) Venn diagram demonstrating the unique and common KEGG pathways identified as significant during ORA enrichment using DEGs from the RNA-seq comparisons (WT compared to clone 1 or clone 4) and DIPs from ATAC-seq comparison (WT compared to clone 4).

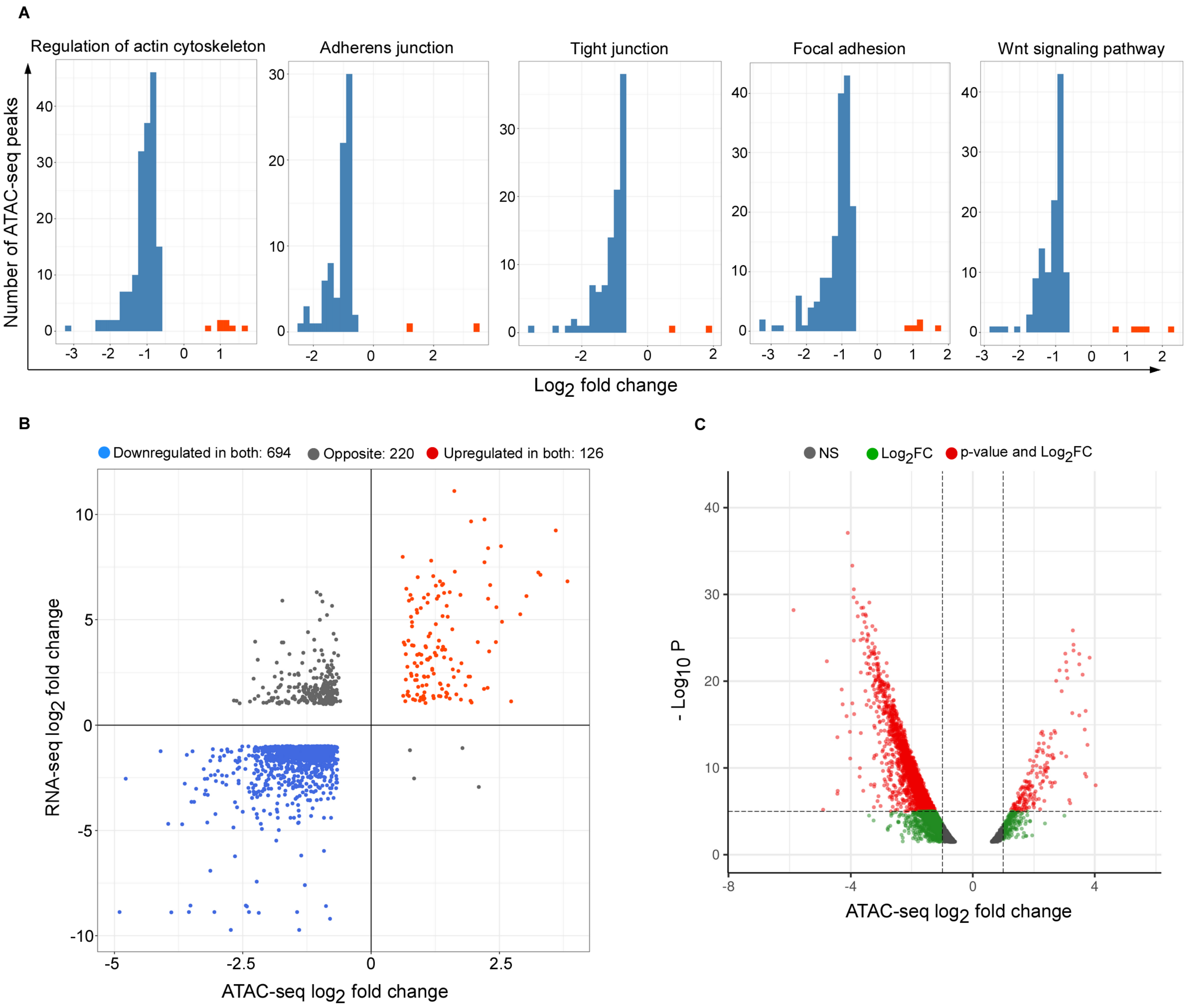
A
Regulation of actin cytoskeleton
Adherens junction
Tight junction
Focal adhesion
Wnt signaling pathway
Number of ATAC-seq peaks
Log2 fold change
B
Downregulated in both: 694
Opposite: 220
Upregulated in both: 126
RNA-seq log2 fold change
ATAC-seq log2 fold change
C
NS
Log2FC
p-value and Log2FC
- Log10 P
ATAC-seq log2 fold change

**Figure S7. Oct4 is predominantly a positive regulator of chromatin landscape in epidermal keratinocytes. Related to Figure 6.**

(A) Bar plots showing the distribution of the significant $log_2$ fold changes in DIPs that overlap genes involved in the five pathways of interest, comparing WT to clone 4 from the ATAC-seq dataset. Red and blue bars correspond to upregulated and downregulated DIPs in clone 4, respectively.

(B) RNA-seq and ATAC-seq datasets comparing WT to clone 4: genes that are both upregulated (DEGs) and located in regions with increased chromatin accessibility (increasing DIPs) are indicated in red, while genes that are both downregulated and located in regions with decreased chromatin accessibility (decreasing DIPs) are highlighted in blue. Dark grey points correspond to genes with DEGs and DIPs with opposite directions (elevated mRNA with decreased chromatin accessibility, or vice versa). Peaks correspond to promoter regions +/- 2kb from TSS. LogFC>1, adj. p-value<0.05.

(C) Volcano plot indicating the adjusted p-values against the $log_2$ fold change of negative (closed chromatin) and positive (opened chromatin) differentially expressed peaks (ATAC-seq dataset) of clone 4 with respect to the WT at gene promoter regions +/- 2kb from TSS.

**Table S1. Amplicon sequencing results. Related to Figures 1, S1.**

| Sample | Mutant Reads | Mutant % | Genotype |
|---|---|---|---|
| WT | 1852 | 0.44 | Homozygous WT |
| Clone 1 | 375526 | 99.87 | Homozygous Mutant |
| Clone 3 | 449927 | 99.85 | Homozygous Mutant |
| Clone 4 | 367516 | 99.86 | Homozygous Mutant |

**Table S2: Primers used for qRT-PCR. Related to Figures 1, 5 and 7.**

| Gene | Sequence 5' → 3' | Application |
|---|---|---|
| *OCT4* Forward | ACATCAAAGCTCTGCAGAAAGAACT | qRT-PCR |
| *OCT4* Reverse | CTGAATACCTTCCCAAATAGAACCC | qRT-PCR |
| *ZO1* Forward | ATTTGTCCGCTCAGCCTGTT | qRT-PCR |
| *ZO1* Reverse | GGCCTCAGAAATCCAGCTTC | qRT-PCR |
| *E-cadherin* Forward | TCTTCAATCCCACCACGTACAA | qRT-PCR |
| *E-cadherin* Reverse | TATTGGGGGCATCAGCATCAG | qRT-PCR |
| *WASP* Forward | GAGAAGCAGAGCCATCCACT | qRT-PCR |
| *WASP* Reverse | GGCAGCAAGTAACTCAGCCA | qRT-PCR |
| *N-WASP* Forward | ACACCAAGCAATTTCCAGCAC | qRT-PCR |
| *N-WASP* Reverse | TGTGCCTCTGAGATTCCACAC | qRT-PCR |
| *Cofilin1* Forward | CTGCAGCGCTCTCGTCTT | qRT-PCR |
| *Cofilin1* Reverse | ACACCGGAGGCCATGTTTC | qRT-PCR |
| *ROCK1* Forward | TGTGGCAATGTGTGAGATGG | qRT-PCR |
| *ROCK1* Reverse | ACAACCCGATTTTCAGCCTTC | qRT-PCR |
| *LIMK1* Forward | TGCATGAGGTTGACGCTACT | qRT-PCR |
| *LIMK1* Reverse | GCAACTCGCTTCCTTCCTCT | qRT-PCR |
| *TSP1* Forward | TGCCTGATGACAAGTTCCAAG | qRT-PCR |
| *TSP1* Reverse | CCAGAGTGGTCTTTCCGCTC | qRT-PCR |
| *Fibronectin* Forward | CACCCCACGCTCAGATACAG | qRT-PCR |
| *Fibronectin* Reverse | ATCACATCCACACGGTAGCC | qRT-PCR |
| *TNC* Forward | AAGCGGGGAATGTTGGGATAG | qRT-PCR |
| *TNC* Reverse | TTAGTCTCCTTTCCACCCCTC | qRT-PCR |
| *CDC42* Forward | ACGCCCCGGTGGAGAA | qRT-PCR |
| *CDC42* Reverse | CATCGCCCACAACAACACAC | qRT-PCR |
| *β-catenin* Forward | AGGACGGTCGGACTCCC | qRT-PCR |
| *β-catenin* Reverse | TCCAACTCCATCAAATCAGCTTG | qRT-PCR |
| *Vimentin* Forward | AGGCGAGGAGAGCAGGATTT | qRT-PCR |
| *Vimentin* Reverse | AGTGGGTATCAACCAGAGGGA | qRT-PCR |
| *β-actin* Forward | CATTCCAAATATGAGATGCGTTGT | qRT-PCR |
| *β-actin* Reverse | GCTATCACCTCCCCTGTGTG | qRT-PCR |
| *GAPDH* Forward | TGCACCACCAACTGCTTAGC | qRT-PCR |
| *GAPDH* Reverse | GGCATGGACTGTGGTCATGAG | qRT-PCR |
